# Golgi-Dependent Copper Homeostasis Sustains Synaptic Development and Mitochondrial Content

**DOI:** 10.1101/2020.05.22.110627

**Authors:** Cortnie Hartwig, Gretchen Macías Méndez, Shatabdi Bhattacharjee, Alysia D. Vrailas-Mortimer, Stephanie A. Zlatic, Amanda A. H. Freeman, Avanti Gokhale, Mafalda Concilli, Christie Sapp Savas, Samantha Rudin-Rush, Laura Palmer, Nicole Shearing, Lindsey Margewich, Jacob McArthy, Savanah Taylor, Blaine Roberts, Vladimir Lupashin, Roman S. Polishchuk, Daniel N. Cox, Ramon A. Jorquera, Victor Faundez

**Affiliations:** Department of Cell Biology, Emory University, Atlanta, Georgia, USA, 30322; Neuroscience Department, Universidad Central del Caribe, Bayamon, Puerto Rico; Neuroscience Institute, Center for Behavioral Neuroscience, Georgia State University, Atlanta, Georgia 30302; School of Biological Sciences Illinois State University, Normal, IL 617901; Center for the Study of Human Health USA, 30322, Emory University, Atlanta, Georgia, USA, 30322; Telethon Institute of Genetics and Medicine (TIGEM), Pozzuoli 80078, Italy; Department of Biochemistry, Emory University, Atlanta, Georgia, USA, 30322; Department of Physiology and Biophysics, University of Arkansas for Medical Sciences, Little Rock, AK, 72205; Institute of Biomedical Sciences, Santiago, Chile

## Abstract

Rare genetic diseases preponderantly affect the nervous system with phenotypes spanning from neurodegeneration to neurodevelopmental disorders. This is the case for both Menkes and Wilson disease, arising from mutations in ATP7A and ATP7B, respectively. The ATP7A and ATP7B proteins localize to the Golgi and regulate copper homeostasis. We demonstrate conserved interactions between ATP7 paralogs with the COG complex, a Golgi complex tether. Disruption of *Drosophila* copper homeostasis by ATP7 tissue-specific transgenic expression caused alterations in epidermis, catecholaminergic, sensory, and motor neurons. Prominent among neuronal phenotypes was a decreased mitochondrial content at synapses, a phenotype that paralleled with alterations of synaptic morphology, transmission, and plasticity. These neuronal and synaptic phenotypes caused by transgenic expression of ATP7 were rescued by downregulation or haploinsufficiency of COG complex subunits. We conclude that the integrity of Golgi-dependent copper homeostasis mechanisms, requiring ATP7 and COG, are necessary to maintain mitochondria functional integrity and localization to synapses.

**Significance Statement:** Menkes and Wilson disease affect copper homeostasis and characteristically afflict the nervous system. However, their molecular neuropathology mechanisms remain mostly unexplored. We demonstrate that copper homeostasis in neurons is maintained by two factors that localize to the Golgi apparatus, ATP7 and the COG complex. Disruption of these mechanisms affect mitochondrial function and localization to synapses as well as neurotransmission and synaptic plasticity. These findings suggest a new principle of interorganelle communication whereby the Golgi apparatus and mitochondria are functionally coupled through homeostatically controlled cellular copper levels.

## Introduction

Rare diseases are genetic maladies that disproportionately affect the nervous system with phenotypes ranging from neurodegeneration to behavioral impairments (Sanders et al., 2019; Lee et al., 2020). The vast collection of mutated human genes and neurological symptomatology offers a wide field for the discovery of novel cellular mechanisms necessary for neuronal function. Here we focus on rare neurological diseases affecting ATP7A, ATP7B, and the COG complex subunit genes, whose products localize to the Golgi complex at steady state (Kaler, 2011; Polishchuk and Lutsenko, 2013; Climer et al., 2015). We report that these ATP7 paralogs and the COG complex converge to maintain copper homeostasis and, unexpectedly, mitochondrial distribution in neurons. Our findings suggest the existence of novel copper-related mechanisms that couple the Golgi complex and mitochondria.

Mutations in ATP7A cause Menkes disease, a systemic copper-depletion affliction caused by intestinal copper malabsorption. Menkes is characterized by childhood systemic and neurological phenotypes. Menkes’ brain phenotypes result from this systemic copper depletion and span from intellectual disability to widespread gray matter neurodegeneration (Menkes, 1988, 1999; Kaler, 2011; Zlatic et al., 2015; Hartwig et al., 2019; Guthrie et al., 2020). In contrast, with the systemic depletion of copper observed in Menkes patients, cell-autonomous ATP7A gene defects cause a cellular overload of copper due to impaired copper efflux from cells (Camakaris et al., 1980; Morgan et al., 2019). This ATP7A cell-autonomous copper phenotype can be recapitulated systemically in Wilson disease. Wilson disease is caused by mutations in the ATP7A paralogue, ATP7B, a copper transporter gene expressed preferentially in the liver. Wilson disease leads to organismal copper overload due to defective copper excretion by the liver. This systemic copper overload causes liver damage, psychiatric symptoms, and lenticular neurodegeneration at late stages of disease (Lutsenko et al., 2007; Kaler, 2011).

ATP7A and ATP7B reside in the Golgi complex at steady state where they contribute to cellular copper homeostasis by sequestering copper into the organelle lumen away from the cytoplasm (Petris et al., 1996; Kaler, 2011; Polishchuk and Lutsenko, 2013). In turn, extracellular copper is taken up by the activity of a plasma membrane copper transporter CTR1 (Kuo et al., 2001; Lutsenko et al., 2007; Kaler, 2011). The expression of ATP7A and CTR1 is controlled by the activity of the COG complex, which tethers vesicles to the Golgi complex (Comstra et al., 2017). The COG complex is an octamer localized to the Golgi apparatus and required for incoming vesicle fusion with the Golgi. Elimination of any one of the eight COG subunits leads to Golgi fragmentation due to destabilization and degradation of the whole octamer (Ungar et al., 2002; Zolov and Lupashin, 2005; Climer et al., 2015; Bailey Blackburn et al., 2016). In addition, we previously demonstrated that cells lacking the COG complex display decreased copper content (Comstra et al., 2017). Human mutations in seven of the eight COG subunits cause a group of diseases collectively known as congenital disorder of glycosylation type II (Wu et al., 2004; Kranz et al., 2007; Foulquier, 2009; Climer et al., 2015). These rare disorders are characterized, in part, by neurodevelopmental pathology and behavioral phenotypes which are loosely similar to neurological phenotypes in Menkes disease. However, whether congenital disorder of glycosylation type II neurological phenotypes may be linked to copper metabolism remains unanswered (Climer et al., 2018).

The convergence of neurological, neurodevelopmental, and psychiatric phenotypes in these three diseases with proteins localizing to the Golgi complex, point to fundamental, neuronal mechanisms necessary to maintain cellular copper homeostasis. These phenotypes raise the question: how are the activities of copper transporters and the COG complex coordinated to maintain copper homeostasis in cells, tissues, and organisms? Here we demonstrate that ATP7 paralogues and the COG complex, both Golgi localized machineries, control synapse development, neurotransmission, and the subcellular localization of mitochondria at synapses. We suggest a new paradigm that the Golgi complex and mitochondria are functionally coupled through copper, an element thus acting as a ‘second messenger’.

## MATERIAL AND METHODS

### Drosophila Strains

The following strains where obtained from the Bloomington Drosophila Stock Center, Bloomington, Illinois: w^1118^ (#5905), pnr-GAL4 (#3039), Ddc-GAL4 (#7009), elav^C155^ -GAL4 (#458), UAS-COG1 RNAi (#38943), UAS-COG5 RNAi (#56882), UAS-COG8 RNAi (#55192), Cog1^e02840^ (#18089), Mito-GFP (#25748). UAS-ATP7-wt was a gift of Richard Burke, Monash University, Australia. The UAS-ATP7 RNAi (#108159) stock was obtained from the Vienna Drosophila Resource Center. GAL4^477^,UAS-mCD8::GFP/CyO,tubP-GAL80;GAL4^ppk.1.9^, UAS-mCD8::GFP (CIV-GAL4 driver), GAL4^477^,UAS-mito-HA-GFP.AP/CyO;ppk::tdTomato, and OregonR (wild-type) were used for dendritic analysis. UAS-mito-HA-GFP.AP line was obtained from Bloomington (#8442).

### Antibodies and Beads

Antibodies used in Drosophila stainings were anti-HRP-FITC (MP Biomedicals, 0855977, RRID:AB_2334736), phalloidin-rhodamine (abcam, ab235138), phalloidin- AlexaFluor 633 (Invitrogen, A22284), anti-Cy3 (Santa Cruz Biotechnology, 166894), the primary antibody rabbit anti-GFP (Sysy 132002, RRID:AB_887725) and the secondary antibody AlexaFluor 488 goat anti-rabbit (Invitrogen).

The following antibodies were used: rabbit anti COG7 from Dr. V. Lupashin’s laboratory, rabbit anti-COG5 (HPA020300, Sigma, RRID:AB_2081406), mouse monoclonal anti-GOLGIN 97 CDF4 (A-21270, Thermo Fisher, RRID:AB_221447), mouse anti-Lamp1 (DSHB H4A3-c, Developmental Studies Hybridoma Bank, RRID:AB_528126), rabbit anti-ATP7B (AB124973, Abcam, RRID:AB_10975785), mouse anti-GAPDH (0411) (sc-47724, Santa Cruz, RRID:AB_627678), mouse anti-myc (c-myc (9E10), sc-40, Santa Cruz, RRID:AB_627268), mouse anti-FLAG M2 (F3165, Sigma, RRID:AB_259529) antibodies. The following beads were used in immunoprecipitations: protein A-Sepharose 4 B Fast Flow from Staphylococcus aureus (P9424, Sigma), Protein G Sepharose 4 Fast Flow (17-0618-01, GE Healthcare), Dynal magnetic beads (110.31, Invitrogen).

### *Drosophila* Immunofluorescence Microscopy

For experiments where larval neuromuscular junctions were analyzed, *Drosophila* stocks were reared in polystyrene vials on standard fly food (900mL milli-Q water, 48g active dry yeast, 120 g cornmeal, 9 g agar, 120 g molasses, 2.4 g tegosept, and 9 mL propionic acid). Animals were housed in a humidified, 25°C incubator (Jeio Tech Co., Inc, IB-15G) with a 12-hour light:dark cycle.

Larval NMJ dissections, immunohistochemistry, and confocal microscopy were performed as described previously (Gokhale et al., 2015b; Mullin et al., 2015; Gokhale et al., 2016). Wandering late third-instar female larvae were dissected with a dorsal incision in a 10 mm cell culture dish in normal HL3 composed of 70 mM NaCl, 5 mM KCl, 21.5 mM MgCl2, 10 mM NaHCO3, 5mM trehalose, 115 mM sucrose, and 5 mM BES, pH 7.2–7.3. Larvae were fixed in 4% paraformaldehyde for 1 hr at room temperature and stained with anti-HRP-FITC (1:500) and anti-phalloidin-rhodamine (1:1000 – data not shown, used for muscle orientation) in PBS supplemented with 0.1% Triton-X 100 (PBS-T) overnight at 4°C. All steps were followed by three 10 min rinses in PBS-T and a final rinse in PBS. The larval preparation was affixed to a poly-L-lysine coated glass side with a drop of Vectashield (Vector Labs H-1000-10) and coverslipped. The coverslip was adhered to the slide with a layer of nail polish in preparation for inverted microscopy. Samples were stored at 4°C until imaged. An inverted Olympus FV1000 microscope was used for confocal imaging of synapses on muscle 6/7 in the A3 segment. Maximum intensity projections were converted to jpegs. Total NMJ length and bouton perimeter measurements were calculated in FIJI (Schindelin et al., 2012). The FIJI Cell Counter plugin was utilized to quantify bouton number (Rai et al., 2018). All files were blind for quantification.

For body wall mosaics, larvae were dissected, fixed, and stained as described above, however, a lateral incision was made to preserve one set of the somatic musculature. The dissection was incubated in the primary antibodies anti-phalloidin-633 (1:500), anti-HRP-Cy3 (1:1000), and rabbit anti-GFP (1:1000) overnight at 4°C and incubated with AlexaFluor 488 goat anti-rabbit (1:1000) with additional anti-phalloidin-633 (1:500), and anti-HRP-Cy3 (1:1000) for 2 hours at room temperature the following day before placement on a slide. Mosaics were imaged using the Multi Area Time Lapse Controller of the Olympus FV1000 confocal microscope. Image stitching was completed, and mosaics converted to jpegs with the Multi Area Time Lapse Viewer of Olympus Fluoview software.

For live confocal imaging, GAL4^477^,UAS-mCD8::GFP;GAL4^ppk.1.9^,UAS-mCD8::GFP virgin flies were crossed to males from individual transgenic fly lines, or outcrossed to *OregonR* males which served as control. Fluorescent neurons from wandering third-instar larvae were imaged on a Zeiss LSM780 confocal microscope as previously described (Das et al., 2017). Briefly, individual animals were placed in a microscopic slide and immersed in 1:5 (v/v) diethyl ether to halocarbon oil and covered with a 22×50 mm coverslip. Images were collected as z-stacks using the 20× dry objective at a step-size of 2 μm. Maximum intensity projections of the images acquired were exported using the Zen blue software. The images were then manually curated using the *Flyboys* software to remove non-specific background noise such as denticle belts. The ImageJ software was used to skeletonize and process these images as previously described (Iyer et al., 2013a; Iyer et al., 2013b), and custom Python algorithms used to compile quantitative data such as total dendritic length and number of branches. For Strahler order analysis, images were reconstructed using NeuronStudio and the branch order extracted using the centripetal branch order function (Rodriguez et al., 2008).

For mitochondrial analysis, virgin female flies from GAL4^477^, UAS-mito-HA.GFP.AP/Cyo;ppk-tdTomato were outcrossed to either males from OregonR (control) or individual transgenic fly lines. Images were acquired as described above and maximum intensity projections of the z stacks were exported using Zen Blue software. Using Adobe Photoshop, images were cropped to a fixed size (6.6 in * 4.5 in) in the same quadrant of the neurons and the mitochondria manually counted along the dendrites in these quadrants. The cropped images were processed as described above to obtain the total dendritic length which was then used to analyze the number of organelles per 1000 pixels dendritic length. For branch point analysis, mitochondria localized at branch points were manually counted and normalized to the total number of branch points.

### *Drosophila* Viability

For experiments measuring animal viability, flies were reared on standard Molasses Food (Genesee Scientific) at 25°C in 12 hr:12hr light:dark cycle. The total number of balanced (control) and unbalanced (experimental) male and female adult offspring from each cross was counted every 1-2 days for 9 days with at least 200 offspring scored. Viability was observed with at least three replicate vials per experiment. 4-5 flies per genotype per sex, where available, were imaged at 4x magnification (Gokhale et al., 2015a; Zlatic et al., 2018).

### *Drosophila* Copper Toxicity

Male and female flies were collected at eclosion and aged 4-7 days. Flies were then starved for 3 hours before being placed in empty vials that contained a filter disc soaked with 200 μl of 100 mM CuSO_4_. ~10 flies per sex per genotype were analyzed with a minimum of 6 replicates. Flies were maintained at 25°C in a 12hr:12hr light:dark cycle and the number of dead flies was assessed daily (Gokhale et al., 2015a; Zlatic et al., 2018).

### *Drosophila* Synaptic Physiology

Drosophila were raised in supplemented fly food (German Formulation, Genesee Scientific), at 20C degrees of temperature and 80% of humidity. Recordings were performed in Modified HL3 solution (10 mM NaHCO3, 5 mM KCl, 4 mM MgCl, 5 mM HEPES, 70 mM NaCl, 5 mM Trehalose, 115 mM Sucrose, pH 7.2). The final Ca^2+^ concentration was set to the desired level indicated in the text. Postsynaptic currents from specific genotypes were recorded at segment A3 of ventral longitudinal muscle 6 as indicated in third-instar larvae using two-electrode voltage clamp with a −80mV holding potential as was previously described (Acharya et al., 2006; Jorquera et al., 2012; Stevens et al., 2012; Astorga et al., 2016). The motor nerves innervating the specified muscles were severed and placed into a suction electrode, so an action potential could be evoked by electrical stimulation at the indicated frequencies using a programmable stimulator (Master8, AMPI). Each stimulus was adjusted to 6V intensity for 0.1ms duration, allowing to recruit both motor neurons and preventing electrotonic induced hyperexcitability.

Data Acquisition and analysis was performed using the National Instrument DAQ system controlled by WinEDR software (University of Strathclyde) on a desktop computer. Electrical recordings were analyzed using Clampfit 10 software (Molecular Devices) or IgorPro 4.0 software (WaveMetrics). Graphs and statistics were generated using OriginPro 9.0 software (OriginLab Corp), and Microsoft Excel routines.

To quantify the post tetanic plasticity in the different genotypes, we adjusted the normalized post-tetanic responses (Fig. 9) with a previously described model that adds depression in the function as P∙e^(−x/τ) +1-D, were P is the post tetanic potentiation (PTP) and D the post tetanic depression (PTD), which decays between two states with a time constant tau. To solve this function, we used numerical solutions by repetitive iteration with solver routines minimizing the summation of the squares of the differences between the data and the model. To help the simulation, we incorporated values of potentiation and depression obtained during the tetanic stimulation. Numerical analysis of normalized post-tetanic enhancement in control and mutant animals generates values for potentiation, depression, and the time constant.

### Quantitation of Total Copper

The measurement of total copper was conducted as previously described (McAllum et al., 2020). Briefly, quantitation of copper metal was performed in three independent collections of 20 third-instar larvae. Copper was measured with an Agilent 7700× series inductively couple plasma mass spectrometer (ICP-MS) using a glass concentric nebulizer (MicroMist, Glass Expansion) and a Scott-type double-pass spray chamber. The instrument was operated in kinetic energy discrimination (KED) mode using helium as a collision gas to remove polyatomic interferences. Quantitation of copper (as m/z 63) was determined with external calibration curves (Accustandard). Yitrium was introduced through a T-piece as an internal standard. The larval samples were digested overnight at room temperature with concentrated (65%) nitric acid (Suprapur^®^grade, Merck) followed by 20 minutes at 90°C. Trace element grade hydrogen peroxide (32%, BDH Aristar^®^ Ultra grade, VWR Analytical) was added to the sample in equal volume to the amount of nitric acid added and then incubated at 70°C for 20 minutes. The sample was then diluted to a final concentration of ~2% nitric acid before analysis.

### Cell Culture

SH-SY5Y (ATCC, RRID:CVCL_0019) and HEK293T (RRID:CVCL_0063) cells were cultured in Dulbecco’s modified Eagle’s medium (DMEM) supplemented with 10% fetal bovine serum (FBS) and 100 μg/ml penicillin and streptomycin at 37°C in 10% CO_2_. HEK293T cells deficient for COG subunits were generated as described (Blackburn and Lupashin, 2016; Willett et al., 2016). HepG2 cells (RRID:CVCL_0027) were grown in Dulbecco’s Modified Eagle’s medium (DMEM) supplemented with 10% FBS (decomplemented at 56°C for 30 min), 2 mM L-glutamine, 100 μg/ml penicillin, and streptomycin at 37°C in 10% CO_2_. HepG2 copper incubations were performed by adding 200μM CuSO_4_ or 400 μM BCS diluted in DMEM at 37°C in a CO2 incubator for 2 hours.

### Immunoprecipitation of ATP7 Paralogues

SH-SY5Y cells were grown to 80-90% confluency in 10 cm tissue culture dishes. On the day of the experiment, cells were washed twice in ice cold PBS followed by lysis in a buffer containing 150 mM NaCl, 10 mM HEPES, 1 mM EGTA, and 0.1 mM MgCl2, with 0.5% Triton X-100 and Complete Protease Inhibitor (Roche 11245200), pH 7.4. The lysate was sonicated and incubated on ice for 30 minutes followed by a clarifying spin at 16,100 × *g* for 10 min. The soluble supernatant was obtained and 500 μg of the protein lysate was added to 30 μl Dynal magnetic beads coated with 5 μl ATP7A antibody (75–142, NeuroMab, RRID: AB_10672736), and incubated for 2 h at 4°C in an end-to-end rotor. As negative controls, immunoprecipitations were done in the presence of a custom made antigenic ATP7A peptide (VSLEEKNATIIYDPKLQTPK, Biosynthesis) prepared in 10 mM MOPS and used at 22 μM (Comstra et al., 2017). Additionally as a non-specific antibody control, we used equivalent amounts of the monoclonal antibody against FLAG (F3165, Sigma, RRID::AB_259529). After the incubation period, magnetic beads were washed 6 times in a buffer containing 150 mM NaCl, 10 mM HEPES, 1 mM EGTA, and 0.1 mM MgCl2, pH 7.4 with 0.1% Triton X-100. Proteins were eluted from the beads with Laemli sample buffer. Samples were resolved by SDS-PAGE and contents analyzed by immunoblot.

For immunoprecipitations using HepG2 cells, cells were solubilized at 4°C for 10 min at room temperature (RT) in a lysis buffer containing 0.5 % Triton X-100, 20 mM Tris/HCl (pH 7.4), 150 mM NaCl, 1mM EDTA (pH: 8), 0.5% NP-40, 10% glycerol, supplemented with 1× protease inhibitor cocktail (Sigma). The mixture was placed into a microfuge tube, kept on ice for 10 min, and then centrifuged at 16,100 × *g* for 15 min at 4°C. Cell lysates were incubated with anti-ATP7B antibody overnight or with anti-FLAG and anti-Myc antibodies as controls. Then protein A or G Sepharose beads were added to each specimen for 4 hours and immune complexes were collected by centrifugation. The beads were then washed and immunoprecipitated proteins were eluted with sample buffer, separated by SDS-PAGE and analyzed by Western blot.

### RNA Interference

Small interfering RNAs (siRNAs) targeting COG5 were purchased from Sigma-Aldrich. The ollowing siRNA were utilized: CCAUUCAUAGUUGGCAGUU and GGACUUUGAAGGAUACUA

Scrambled siRNAs were used as a control. HepG2 cells were transfected with siRNAs using Dharmafect4 (Dharmacon T-2004, Pittsburgh, USA), according to manufacturer instructions.

### RNA Preparation and qRT-PCR

Control cells and COG5-silenced cells were lyzed and total RNA purified by QIAshredder (Qiagen) and extracted with RNeasy Protect Mini Kit (Qiagen) using standard conditions. Total RNA (1 μg) was reverse-transcribed by QuantiTect Reverse Transcription kit (Qiagen) according to the manufacturer’s instructions. qRT-PCR experiments were performed using Light Cycler 480 Syber Green MasterMix (Roche) for cDNA amplification and LightCycler 480 II (Roche) for signal detection. qRT-PCR results were analyzed using the comparative Ct method normalized against housekeeping gene β-Actin. The following primers were used:

β-ACTIN forward (5’-AAGAGCTACGAGCTGCCTGA-3’), β-ACTIN reverse (5’-GACTCCATGCCCAGGAAGG-3’), COG5 forward (5’-AGAGCCCGACTTGAAGTGGAAA-3’), COG5 reverse (5’- ACCTGAAGAGCTGTTCCGACTT-3’).

### Immunofluorescence

Control and COG5-silenced cells were fixed (10 min, 4% paraformaldehyde in 0.2 M HEPES) directly or 2h after incubation with 200μM CuSO_4_. Fixed cells were then incubated with blocking/permeabilizing solution (0.5% bovine serum albumin, 0.1% saponin, 50 mM NH4Cl in PBS) for 20-30 min. Primary and secondary antibodies were diluted in blocking/permeabilizing solution and added to the cells for 1h and 45 min, respectively. Samples were examined with a Zeiss LSM 700 confocal microscope equipped with a 63X 1.4 NA oil objective.

### Extracellular Flux Analysis of Tissue Culture Cells

All oxygen consumption rates were measured on the Seahorse XFe96 Analyzer (Seahorse Bioscience) following manufacturer recommendations. XFe96 extracellular flux assay kit probes (Seahorse Bioscience 102601-100) were incubated with the included calibration solution overnight at 37°C under non-CO_2_ injected conditions. Cells were trypsinized, counted (BioRad TC20 automated Cell Counter) and seeded into XFe96 cell culture microplates (Seahorse Bioscience 101085-004) at 20,000 cells/well and incubated at 37°C with 10% CO_2_ in complete culture media for 24 hours before initialization of the stress test. The next day, wells were washed twice in Seahorse stress test media consisting of Seahorse XF base media (Seahorse Bioscience 102353-100), 2mM L-Glutamine (Hyclone SH30034.01), 1mM sodium pyruvate (Sigma S8636), and 10mM D-glucose (Sigma G8769) pH7.4 before final volume added and then incubated at 37°C in non-CO_2_ injected conditions for 1 hour prior to stress test. Seahorse injection ports were filled with 10-fold concentrated solution of disulfiram (Sigma 86720), oligomycin A (Sigma 75351), carbonyl cyanide-4-(trifluoromethoxy)phenylhydrazone (FCCP) (Sigma C2920), and rotenone (Sigma R8875)/Antimycin A (Sigma A8674) for final testing conditions of disulfiram (0.1 μM), oligomycin (1.0 μM), FCCP (0.125 μM), rotenone (0.5 μM), and antimycin A (0.5 μM). The flux analyzer protocol conditions consisted of 3 basal read cycles, 40 reads following disulfiram injection, and 3 reads following each injection of oligomycin A, FCCP, and rotenone plus antimycin A. Each read cycle consisted of a 3 minutes mix cycle, followed by a three minute read cycle where oxygen and pH levels were determined over time. The Seahorse Wave Software version 2.2.0.276 was used for data analysis of oxygen consumption rates. Experiments were repeated in quadruplicate and aliquots of each batch of stress test media were sent for inherent copper concentration analysis by ICP mass spectrometry performed by the Center for Applied Isotope Studies (CAIS) at the University of Georgia. Non-mitochondrial respiration was determined as the lowest oxygen consumption rate following injection of rotenone plus antimycin A. Basal respiration was calculated from the oxygen consumption rate just prior to oligomycin injection minus the non-mitochondrial respiration. Acute response was calculated from the difference in oxygen consumption rates just prior to disulfiram injection to just prior to oligomycin injection. Oligomycin A sensitivity was calculated as the difference in oxygen consumption rates just prior to oligomycin injection to the minimum oxygen consumption rate following oligomycin injection but prior to FCCP injection.

### Evolutionary Rate Covariation Analysis

We used the web engine https://csb.pitt.edu/erc_analysis/ (Clark et al., 2012, 2013; Findlay et al., 2014) using as inputs the CORUM annotated complexes database (Ruepp et al., 2010).

### Statistical analysis

Experimental conditions were compared using Synergy Kaleida-Graph, (version 4.1.3)) or Aabel NG2 v5 x64 (Gigawiz) as specified in each figure.

## Results

### P-ATPase Copper Transporters Interact with the COG Complex

ATP7A and ATP7B are paralogs present in chordates, which originated from a single ATP7 gene found in the last common ancestor to eumetazoans. The *Drosophila* genus retained this single ATP7 gene (http://www.treefam.org/family/TF300460). We previously demonstrated that human ATP7A and the COG complex biochemically interact in cells (Comstra et al., 2017). Moreover, mutations of COG complex subunits reduced the expression of ATP7A and CTR1, resulting in cellular copper depletion (Comstra et al., 2017). As ATP7B shares 52.5% identity with ATP7A (alignment of Uniprot entries ATP7A_HUMAN and ATP7B_HUMAN), then ATP7B may also interact with the COG complex. To test this hypothesis, we immunoprecipitated endogenous ATP7A or ATP7B from neuroblastoma or hepatoma human cell lines, respectively, and assessed the presence of COG complex subunits by immunoblot with antibodies against COG5 and COG7 (Fig. 1). Both ATP7A and ATP7B immunoprecipitated COG complex subunits (Fig. 1A lane 4 and 1B lanes 3 and 4). The selectivity of these ATP7A/B coprecipitations with COG subunits was determined with control immunoprecipitations with unrelated antibodies (Fig. 1A lane 3 and 1B lanes 3’ and 4’), out-competing immune complexes with the ATP7A antigenic peptide (Fig. 1A lane 5), or by blotting ATP7B immunoprecipitates with GAPDH antibodies (Fig. 1B). Similar to ATP7A, the association of ATP7B with the COG complex remained after challenging cells with excess extracellular copper (Fig. 1B, compare lanes 3 and 4). Downregulation of COG5 in hepatoma cells (Fig. 1C) disrupted Golgi morphology and decreased ATP7B expression, both at Golgi and lysosomal compartments, as revealed by co-labeling cells with ATP7B and GOLGIN95 or LAMP1 antibodies, respectively (Fig. 1D). These findings demonstrate a conserved interaction between the COG complex with ATP7 paralogs. Moreover, these results establish downregulation of copper transporters as a conserved phenotype after disruption of COG complex function.

**Fig. 1.**
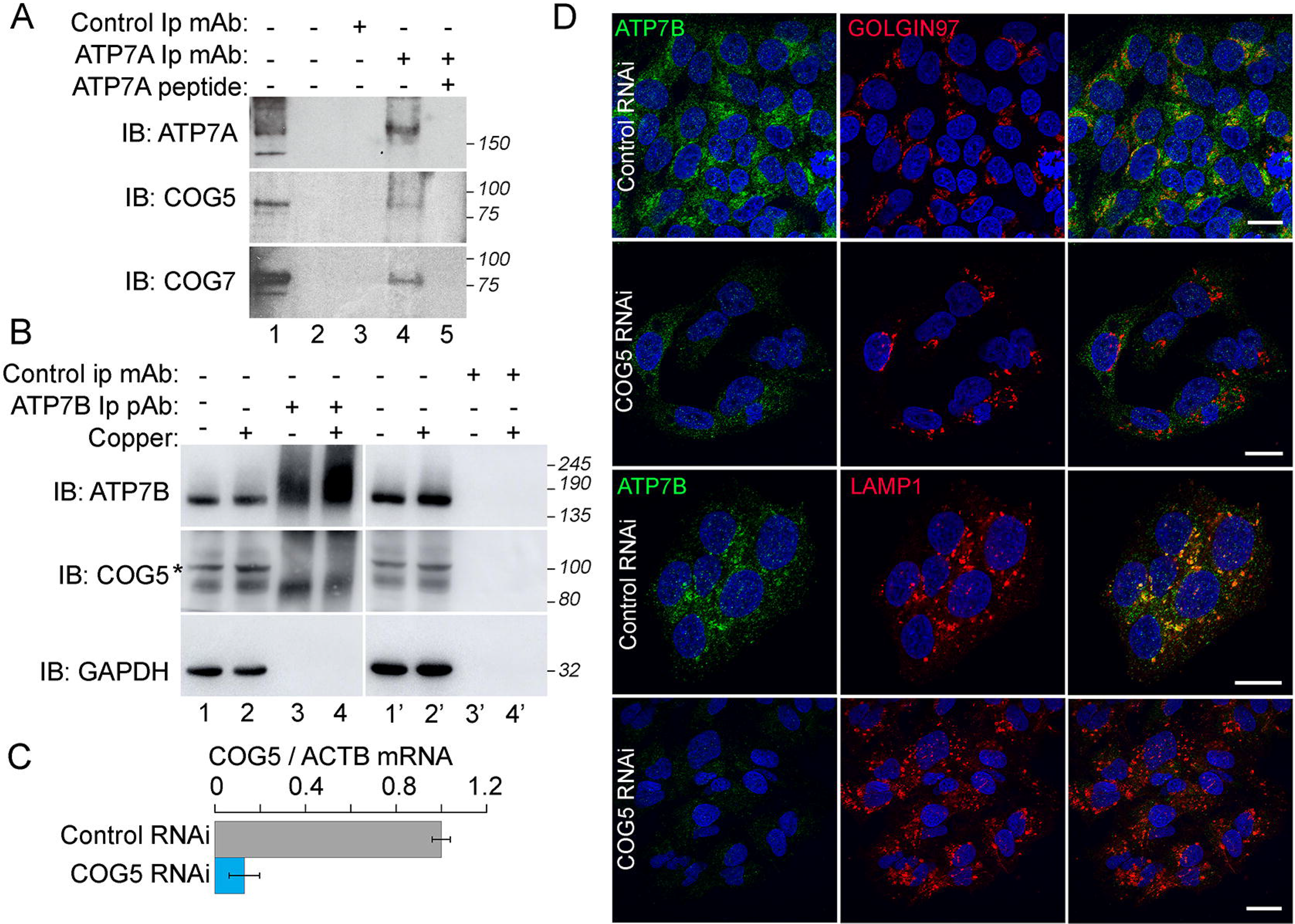
Biochemical and Genetic Interactions between COG Complex Subunits and ATP7 Paralogs. A) SH-5YSY and B) HepG2 cells were incubated in normal media or media supplemented with a 200 μM copper excess (Fig. 1B, even lanes). Detergent soluble extracts from A) SH-5YSY and B) HepG2 cells were immunoprecipitated with antibodies against ATP7A or ATP7B, respectively. The presence of COG complex in the immunoprecipitates was detected with antibodies against COG5 and COG7. Controls were performed with unrelated antibodies (Fig. 1A, lane 3 or Fig. 1B, lanes 3’-4’) or with the ATP7A antigenic peptide (Fig. 1A, lane 5). Asterisk marks non-specific band. A0 Lane 1 corresponds to input. B) Lanes 1,2,1’ and 2’ correspond to inputs. C) siRNA knockdown of COG5 in HepG2 cells. Relative expression corresponds to the ratio between COG5 and ACTB (n=3). D) Immunomicroscopy analysis of the expression and localization of ATP7B in Golgi (untreated cells) and lysosomes (CuSO_4_-treated cells). These compartments were detected with antibodies against GOLGIN97 and LAMP1, respectively. Bar 10 μm.

### Synaptic Development Requires ATP7- and COG-Dependent Copper Homeostasis

We tested the phylogenetic conservation of the interactions between the COG complex and copper transporters in adult and developing *Drosophila* neurons. We took advantage of the fact that the expression of mammalian ATP7A, ATP7B, and CTR1 is reduced in cells that lack COG complex subunits (Fig. 1 and (Comstra et al., 2017)). We reasoned that *Drosophila* neurons would help us to genetically test whether the COG complex balances the expression of *Drosophila* ATP7 and CTR1 to control copper homeostasis. We analyzed the COG complex function balancing the expression of these copper transporters in developing *Drosophila* third-instar larvae motor neurons and cuticle sensory neurons. We reasoned that cell autonomous and systemic changes in transporter expression should differentially and predictably affect copper content in developing tissues and animals (Fig. 2A). For example, cell autonomous ATP7 overexpression would decrease cellular copper, a model of copper efflux (Fig. 2A) (Norgate et al., 2006; Burke et al., 2008; Binks et al., 2010). In contrast, copper content after systemic ATP7 overexpression would be the result of increased midgut copper absorption plus a decreased cellular copper content in internal tissues (Fig. 2A)(Bahadorani et al., 2010). Conversely, cell autonomous ATP7 knockdown increases cellular copper, much like in Menkes disease cultured cells (Fig. 2A)(Norgate et al., 2006; Burke et al., 2008; Binks et al., 2010). Yet, systemic ATP7 inhibition would be dominated by midgut copper absorption defects, thus mimicking Menkes disease (Fig. 2A).

**Fig. 2.**
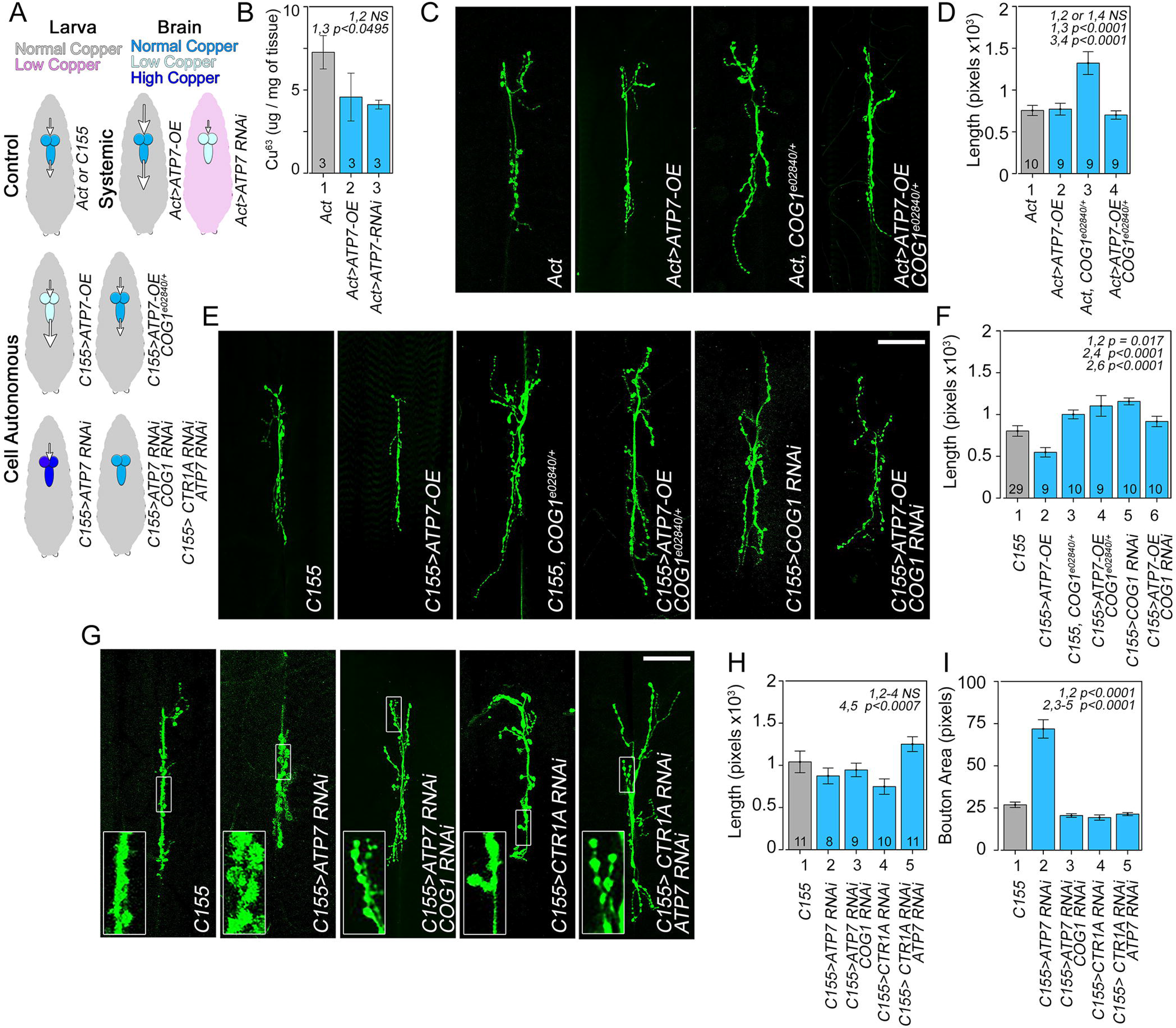
Copper Homeostasis by the COG complex and Copper Transporters Modulate Glutamatergic Synapse Morphology. A) Model of predicted larval and brain copper status in different genotypes. Size of the arrow depicts the magnitude of the influx or efflux of copper into the brain. Note that changes in copper content in brain and whole larvae can be independently modulated by neuronal cell autonomous (C155>) or systemic (Actin driver, Act>) modifications of ATP7 gene expression. B) ICP mass spectrometry determinations of total larval copper content in UAS-ATP7 or UAS-ATP7 RNAi transgenes expressed from a ubiquitous Actin-GAL4 driver. N= 3 independent batches of 20 third-instar larvae each. p value was calculated with two-tailed Mann-Whitney rank-sum test. Muscle 6-7 neuromuscular junctions from wild type animals and animals expressing ATP7, COG and CTR1 UAS-transgenes stained with anti-HRP antibodies (C, E, and G). C) Synapses from animals carrying the UAS-ATP7 transgene expressed under the systemic driver Actin-GAL4, the *COG1*^*e02840/+*^ genotype, each in isolation or their combination. Bar 50 μm. D) Depicted total branch length quantifications of animals in C. E) Synapses from animals carrying the UAS-ATP7 transgene, UAS-COG1 RNAi expressed under the C155-GAL4 driver, or the *COG1*^*e02840/+*^ genotype, each in isolation or their combinations. F) Depicted quantifications of animals in (E). G) Synapses from animals carrying the UAS-ATP7 RNAi, UAS-CTR1 RNAi expressed with the C155-GAL4 driver, or their combinations. H-I) Shows total branch length and bouton size area quantifications, respectively, of animals in G. Number of animals per genotype (n) are presented by the number at the base each column. One Way ANOVA followed by two-tailed Fisher’s Least Significant Difference Comparison.

We first tested these predictions by systemic overexpression of ATP7 or downregulation of ATP7 in all tissues using the Actin-GAL4 driver (Fig. 2A). Systemic overexpression of ATP7 did not significantly modified the larval copper content (Fig. 2B). In contrast, ubiquitously expressed ATP7 RNAi transgene partially decreased the copper content in whole larvae, partially mimicking the organismal copper depletion phenotype observed in Menkes disease (Fig. 2B). To determine if systemic ATP7 overexpression affects synaptic structure, we analyzed neuromuscular junction morphology and found that overexpression of ATP7 had no effect on synapse size (Fig. 2C-D). In contrast, flies heterozygous for COG1 had increased synapse size measured as the total length of all branches (Fig. 2C-D, *COG1*^*e02840/+*^), a phenotype rescued by overexpression of ATP7 (Fig. 2C-D). This result suggests that the effects of reduced COG1 are due in part to the COG effects on endogenous ATP7 expression (Fig. 2C-D). If loss of COG1 results in reduced ATP7 expression, then phenotypes caused by neuronal cell autonomous ATP7 overexpression should be reverted by either systemic or neuronal-selective downregulation of the COG1 subunit. We find that neuronal-specific overexpression of ATP7 using C155-GAL4 results in decreased synapse size (Fig. 2E-F) and this phenotype was rescued by either COG1 heterozygosity or neuronal-specific knockdown of COG1 (Fig. 2E-F). We conclude that COG1 and ATP7 interact genetically to modulate synapse morphology. Second, we manipulated the expression of CTR1 and the COG complex cell autonomously in larval motor neurons in which ATP7 expression is downregulated (Fig. 2A, G-I). Neuron-specific downregulation of ATP7 did not change the neuromuscular junction size (Fig. 2G-H) but increased the size of synaptic boutons (Fig. 2G and I, see insert). This phenotype was rescued by either neuronal-specific COG1 or CTR1 knockdown, indicating that COG1 downregulation modulates the entry of copper from the plasma membrane phenocopying CTR1 downregulation (Fig. 2G and I). These results demonstrate that the COG complex regulates cellular copper homeostasis by controlling the function of ATP7 and CTR1 in developing motor neurons.

To assess whether ATP7 and COG genetic interactions modulating synapses were just restricted to developing motor neurons, we measured dendritic arborization complexity in *Drosophila* third-instar larvae class IV (C-IV) dendritic arborization (da) epidermal sensory neurons (Fig. 3). The C-IV da neuron possesses an elaborated and stereotypic dendritic arbor whose complexity is responsive to gene perturbations (Iyer et al., 2013a; Singhania and Grueber, 2014; Das et al., 2017). We overexpressed ATP7 in a cell-autonomous manner, a model of copper efflux, using the ppk-GAL4 driver (Fig. 2A)(Yang et al., 2009). ATP7 expression significantly decreased the complexity of the dendritic arbor, as determined by the total dendritic length and number of branches (Fig. 3A-C). COG1 haploinsufficiency with the *COG1*^*e02840/+*^ allele did not significantly change dendritic arbors (Fig. 3A-C). However, this COG1 gene dosage reduction was enough for suppressing the ATP7 overexpression dendritic phenotype (Fig. 3A-C). The dendritic phenotype caused by ATP7 overexpression and its rescue were more prominent in terminal branches, as indicated by the reversed Strahler analysis, in which distal branches of the 1 and 2 categories were the most affected as compared to branches closer to the cell body (Fig. 3D). These results demonstrate that the COG complex balances the expression of copper influx and efflux transport activities from cells. In turn, these changes in copper homeostasis modulate dendritic and presynaptic morphology in diverse developing neuronal types.

**Fig. 3.**
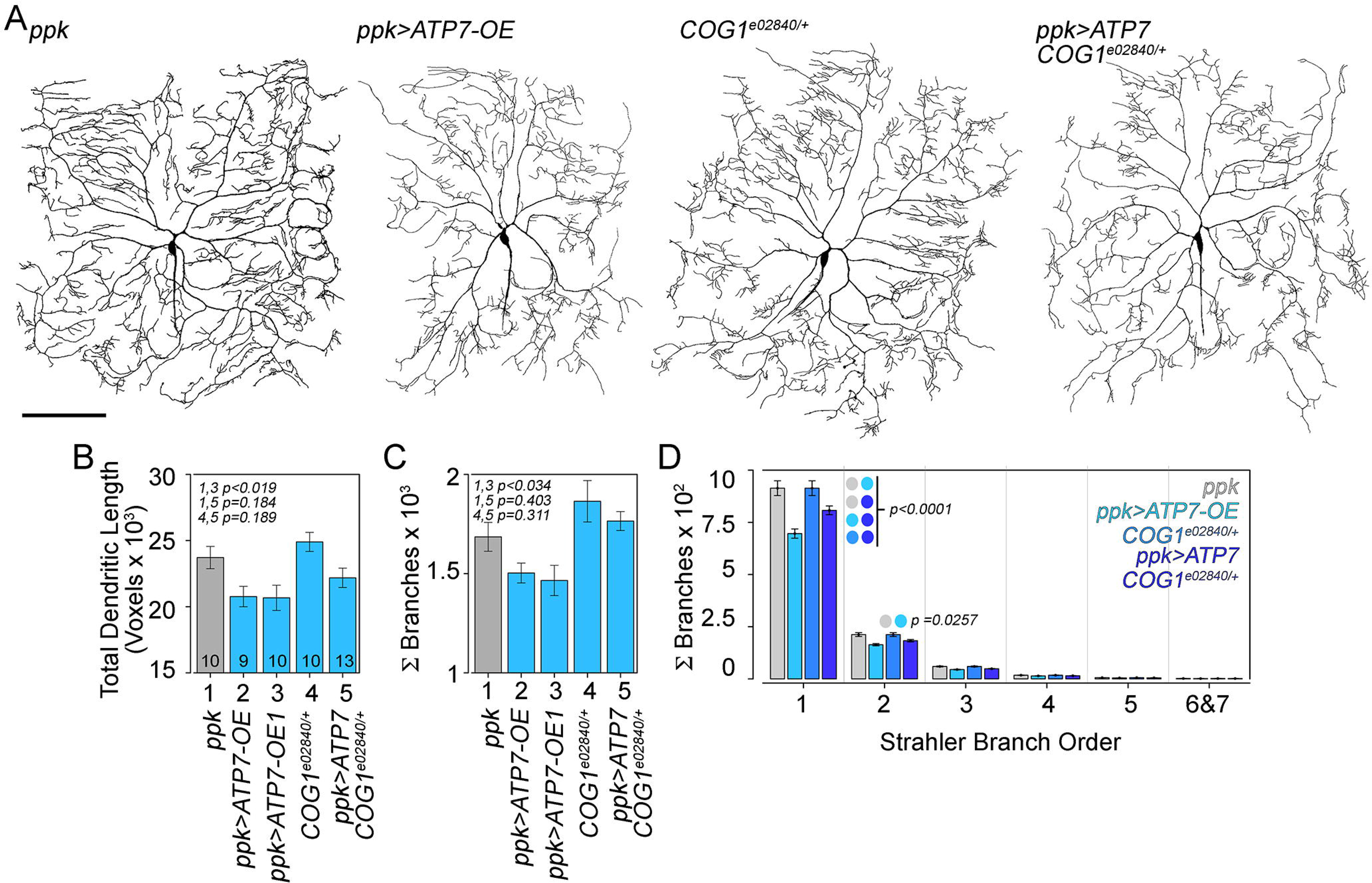
The COG Complex and ATP7 Modulate Sensory C-IV da Neuron Dendrite Architecture. A) Representative live confocal images of C-IV da neurons of the specified genotypes labeled by GFP driven by CIV-GAL4 (477;ppk). B-D show quantitative analysis of total dendritic length and total number of branches in the specified genotypes. 200 μm. D) Depicts reversed Strahler branch order analysis. Terminal branches correspond to Strahler number 1 and 2. Average ± SEM. n of animals per genotype are presented by the number at the base each column. For B-C comparisons were made with One Way ANOVA followed by two-tailed Fisher’s Least Significant Difference Comparison. D, analysis was made with Two-way ANOVA followed by Tukey’s multiple comparisons test.

### The COG Complex Genetically Interacts with ATP7 Transporters in Adult *Drosophila* Tissues

We applied the same experimental logic utilized for larval synapses to adult *Drosophila* catecholaminergic neurons. These cells are only sensitive to ATP7 expression manipulation when the animals are exposed to excess dietary copper (Comstra et al., 2017). Moreover, copper-dependent molecules and transporters are highly-expressed in vertebrate catecholaminergic cells (Xiao et al., 2018). As we previously demonstrated, copper feeding control animals increased mortality close to a 100% (Fig. 4A, column *1*) (Comstra et al., 2017). Catecholaminergic neuron ATP7 overexpression using the Ddc-GAL4 driver decreased copper-induced mortality close to 30% and 60% in adult females and males, respectively (Fig. 4A, columns *1* and *2*). We interpret this protective effect as an increased copper efflux due to overexpressed ATP7 (Fig. 2A). A similar protective outcome was achieved by downregulation of the COG1, COG5, or COG8 subunits (Fig. 4A, column *1* compare to *3*, *5*, and *7*). Here, the protective effect of COG downregulation likely uncovers a decreased CTR1 activity necessary for copper influx into catecholaminergic neurons (Fig. 2A). To define how the COG complex balances the expression of ATP7 and a CTR1 activity, we downregulated COG complex subunits in the ATP7 overexpression background and find that this leads to increased mortality as compared to the COG subunit RNAi alone (Fig. 4A, compare columns *3-4*, and *7-8*), suggesting that the COG complex preferentially affects the expression of ATP7 over a copper uptake transporter activity (Fig. 2A). If the COG complex has a stronger interaction with ATP7 than CTR1 in catecholaminergic neurons, then knockdown of ATP7, a model of copper influx, should not protect animals after a copper feeding challenge. Additionally, the ATP7 knockdown phenotype should be suppressed by inhibition of the COG complex due to disruption of copper influx mechanisms (Fig. 2A)(Comstra et al., 2017). As predicted, inhibition of ATP7 mortality reach near 100% after copper feeding and was indistinguishable from wild type animals (Fig. 4B, compare columns *1* and *2*). Moreover, the effect of copper feeding on mortality were suppressed by inhibition of the COG subunits alone or in combination with inhibition of ATP7 (Fig. 4B, compare columns *3-4*, *5-6*, and *7-8*). These results demonstrate that in neurons expressing ATP7, COG complex deficiency preferentially suppresses the effects of ATP7 overexpression (Fig. 2A). These findings demonstrate the COG complex balances cellular copper homeostasis by modulating the expression of ATP7 and copper influx activity, presumably through CTR1.

**Fig. 4.**
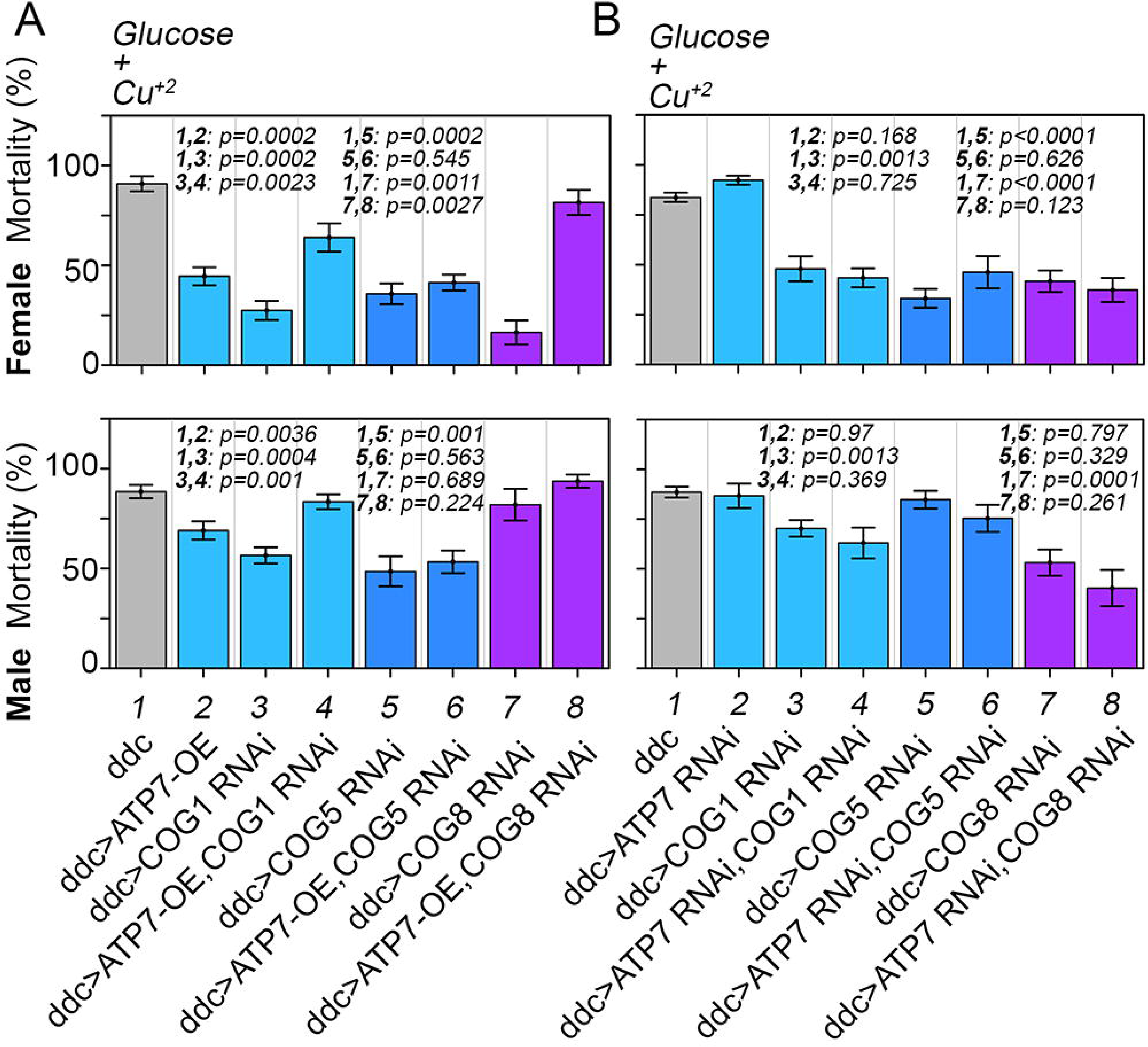
Interactions between ATP7 and the COG Complex are Conserved in *Drosophila* Adult Dopaminergic Neurons. Male and female adult animals expressing UAS-ATP7A (A), UAS-ATP7 RNAi (B), or RNAi against COG1, COG5, or COG8 alone or in combination in dopaminergic neurons using the ddc-GAL4 driver (A-B). Mortality was induced by copper feeding for 2-3 days. For experiments assessing UAS-ATP7 in both sexes, we used 10 experiments with 86-96 animals in total per genotype and sex, except for UAS-COG8 RNAi experiments were n=7-8 with 46-77 animals used in total. For experiments assessing UAS-ATP7 RNAi in both sexes, we used 12-38 experiments with 80-336 animals in total per genotype and sex. p values calculated with Kruskal-Wallis test followed by pairwise comparisons with two-tailed Mann-Whitney rank-sum test. See Extended Data Fig. 4-1

We validated these genetic interactions and their interpretations in the *Drosophila* adult epidermis where the interplay of ATP7 and CTR1 in controlling copper homeostasis has been precisely established. In this tissue, overexpression of ATP7 with the epidermal pnr-GAL4 driver depletes cellular copper, a phenomenon rescued by increased expression of the plasma membrane copper uptake transporter CTR1 (Norgate et al., 2006; Burke et al., 2008; Binks et al., 2010; Hwang et al., 2014; Mercer et al., 2016). We reasoned that, in contrast with neurons, phenotypes caused by epidermal overexpressed ATP7 would be additive with epidermal COG complex subunit RNAi. Thus, supporting the concept that copper homeostasis is chiefly controlled through a CTR1 activity in the epidermis (Fig. 4-1). Overexpression of ATP7 caused incomplete dorsal closure and depigmentation of the thorax and abdomen of adult female and males flies (Fig. 4-1A). These phenotypes were maladaptive, as demonstrated by decreased animal survival to ~38% and ~65% in adult females and males, respectively (Fig. 4-1B). These ATP7-dependent epidermal phenotypes were absent in animals where the expression of the COG complex was reduced by RNAi against any one of three COG subunits (Fig. 4-1B). However, simultaneous expression of ATP7 in animals where the COG complex was downregulated by RNAi significantly decreased survival to ~22 and ~35% in adult females and males, respectively (Fig. 4-1B). This doubling in mortality represents an additive phenotype. These results indicate that the COG complex modulates copper homeostasis by preferentially controlling the expression of a CTR1 activity in epidermal cells (Fig. 4-1C). These epidermal findings further validate our genetic strategy to examine whether an intact COG complex is requirement for balancing copper influx and efflux in neurons.

### Mitochondrial Respiration Requires a COG- and Copper-Dependent Mechanisms

A possible target of copper dyshomeostasis in motor and sensory *Drosophila* neurons is the mitochondria as both genetic defects in ATP7A and ATP7B perturb mitochondrial morphology and function (Yoshimura and Kudo, 1983; Yamano and Suzuki, 1985; Onaga et al., 1987; Yamano et al., 1988; Gu et al., 2000; Zischka et al., 2011; Bhattacharjee et al., 2016; Lichtmannegger et al., 2016; Guthrie et al., 2020). Similarly, COG complex mutants decrease tetrazolium salt metabolization mediated by the NAD(P)H-dependent oxidoreductase and dehydrogenase enzymes, which localize both to the cytoplasm and mitochondria (Berridge et al., 2005; Comstra et al., 2017). This suggests that the COG complex could modulate mitochondrial function in a copper-dependent manner. We tested this hypothesis using phylogenetic-bioinformatic analysis and measuring mitochondrial respiration with Seahorse^™^ oximetry as well.

We used evolutionary rate covariation to determine functional associations between either the ATP7A and ATP7B proteins or the COG complex with subunits of each one of the five mitochondrial respiratory chain complexes. This approach allows us to examine coevolutionary forces that act on genes even when their products do not physically interact or may reside in different compartments (Fig. 5A-B, ERC) (Clark et al., 2012, 2013; Findlay et al., 2014). The phylogenic correlation between ATP7A and ATP7B was significant with mitochondrial complexes I, III, and IV, the latter, a copper-containing respiratory chain complex (Fig. 5A) (Horn and Barrientos, 2008). Moreover, the eight subunits of the COG complex also covaried significantly with complex I (Fig. 5A). We analyzed the evolutionary rate covariation among all pairs of a gene group integrated by ATP7A, ATP7B, the COG complex, and the mitochondrial complex I subunits. Gene group analysis of the evolutionary rate covariation among these 48 genes indicated that this gene set is significantly correlated phylogenetically (Fig. 5B, p=0.00130). Also, pairwise analyses showed that ATP7A coevolved with ATP7B (ERC scores >0.3, p<0.05), and multiple complex I subunits (Fig. 5B-C). Conversely, COG complex subunits significantly coevolved with ATP7B, and complex I subunits (Fig. 5B-C). These results support a model where ATP7A, ATP7B, and the COG complex participate in a pathway shared with mitochondria.

**Fig. 5.**
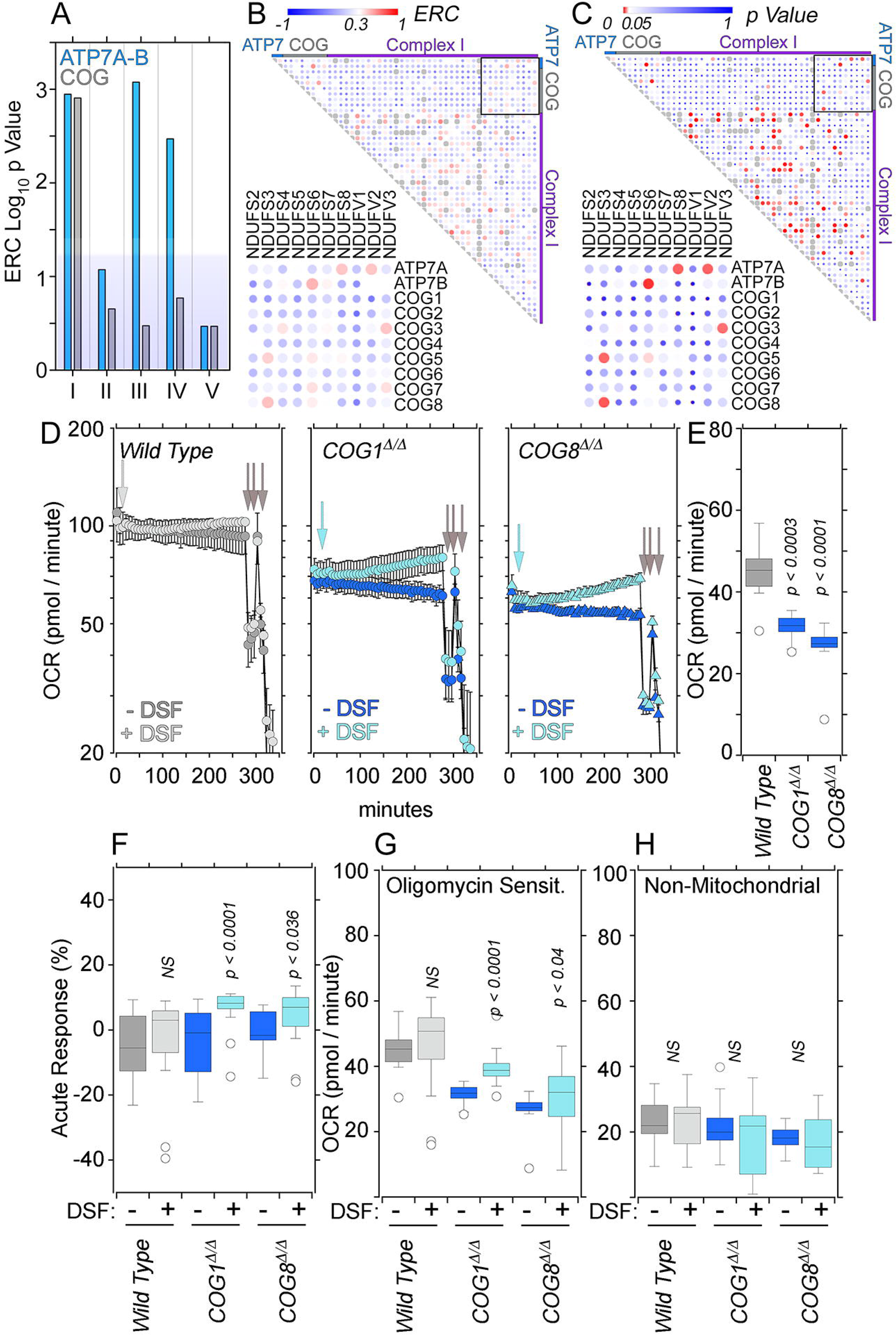
The COG Complex Activity is Required for Mitochondrial Respiration. (A-C) Evolutionary rate covariation between gene groups constituted by human ATP7A and B or the COG complex subunits with subunits of the five mitochondrial respiratory complexes defined by the CORUM database. A) Depicts the group probability of evolutionary covariation. B) Presents evolutionary rate covariation (ERC) for pairs of genes in a gene group that includes ATP7A and B, the eight COG complex subunits, and the 38 nuclear encoded subunits of the mitochondrial respiratory complex I (CORUM complex ID:388). ERC threshold >0.3 identifies covariated gene pairs. C depicts p values for genes pairs presented in B. alpha <0.05 Wilcoxon rank sum test against 100,000 permutations. ERC and p values are represented by circle color and size. D to H) COG null cells have defective respiration that can be ameliorated with a copper ionophore. Extracellular Seahorse flow oxymetry was used to measure oxygen consumption rate (OCR) in wild type, COG1^Δ/Δ^, and COG8^Δ/Δ^ HEK293 cells with or without the addition of 100 nM disulfiram in the presence of 180 nM copper. (D) Depicts (OCR) each time points over 300 minutes, while (E) depicts the average basal OCR, and (F) shows the acute response. Mitochondrial respiration was isolated by the addition of 0.1μM oligomycin (G), and non-mitochondrial respiration was measured by the addition of 0.5μM rotenone and antimycin A(H). N=4, non-parametric Kruskal Wallis test followed by pairwise Mann-Whitney U test comparisons.

We tested whether COG genetic defects could impair mitochondrial function by measuring oxygen consumption rates with Seahorse^™^ technology (Divakaruni et al., 2014). We used HEK293 cells where both genomic copies of the COG subunits 1 (COG1^Δ/Δ^) or 8 (COG8^Δ/Δ^) genes were edited by CRISPR-Cas9 (Bailey Blackburn et al., 2016; Blackburn and Lupashin, 2016). These COG null cells phenocopy established COG complex deficiency phenotypes, which include defective Golgi-dependent glycosylation of membrane proteins and their degradation in lysosomal compartments (Shestakova et al., 2006; Bailey Blackburn et al., 2016; Blackburn and Lupashin, 2016). COG null HEK293 cells had a lower basal oxygen consumption rate compared to wild type controls (Fig. 5D-E). To determine if this reduced basal respiration is due to impaired copper availability we treated cells with disulphiram, a copper ionophore (Allensworth et al., 2015). We find that the basal respiration rate in wild type cells in the presence of 180 nM copper was not changed even after 4 hours (Fig. 5D and F, acute response). In contrast, the addition of copper-disulfiram to COG1^Δ/Δ^ and COG8^Δ/Δ^ HEK293 cells significantly improved the basal respiration rate, a response that was evident at 2 hours and progressively increased up to 4 hours of incubation with copper-ionophore (Fig. 5D and F, acute response). This selective improvement in oxygen consumption by COG null cells was sensitive to the mitochondrial ATPase inhibitor oligomycin (Divakaruni et al., 2014), thus demonstrating the mitochondrial origin of the copper-disulfiram rescue of the respiration phenotype in COG-null HEK293 cells (Fig. 5D and G). Non-mitochondrial sources of oxygen consumption were not affected by genotype or copper ionophore addition, as determined by measuring oxygen consumption in the presence of antimycin and rotenone, inhibitors of cytochrome C reductase and complex I, respectively (Fig. 5D and H) (Divakaruni et al., 2014). These results demonstrate that mitochondrial respiration defects observed in COG deficient cells can be acutely corrected by circumventing impaired copper transport with a copper ionophore. These phylogenetic and functional results show that COG complex-dependent copper homeostasis is necessary for mitochondrial function.

### ATP7- and COG-Dependent Copper Homeostasis Sustain Mitochondrial Synaptic Content

We sought to answer whether ATP7- and COG-dependent copper homeostasis was required for normal *Drosophila* mitochondria at motor neuron synapses. To address this question, we generated animals carrying both UAS-ATP7 and a UAS-mitochondria-targeted GFP reporter (Fig. 6, MitoGFP), and the C155-Gal-4 neuronal driver. We focused on the neuronal ATP7 overexpression/copper efflux paradigm because ATP7 overexpression synaptic phenotypes are robustly rescued just by a single gene copy loss of COG1 (Fig. 2E and F). We measured synapse morphology on neuromuscular junctions double-stained with antibodies against HRP and GFP to label the neuronal plasma membrane and mitochondria, respectively. Mitochondria were present in neuromuscular junctions, including muscles 6 and 7, a standard synaptic model (Fig. 6A and B).

**Fig. 6.**
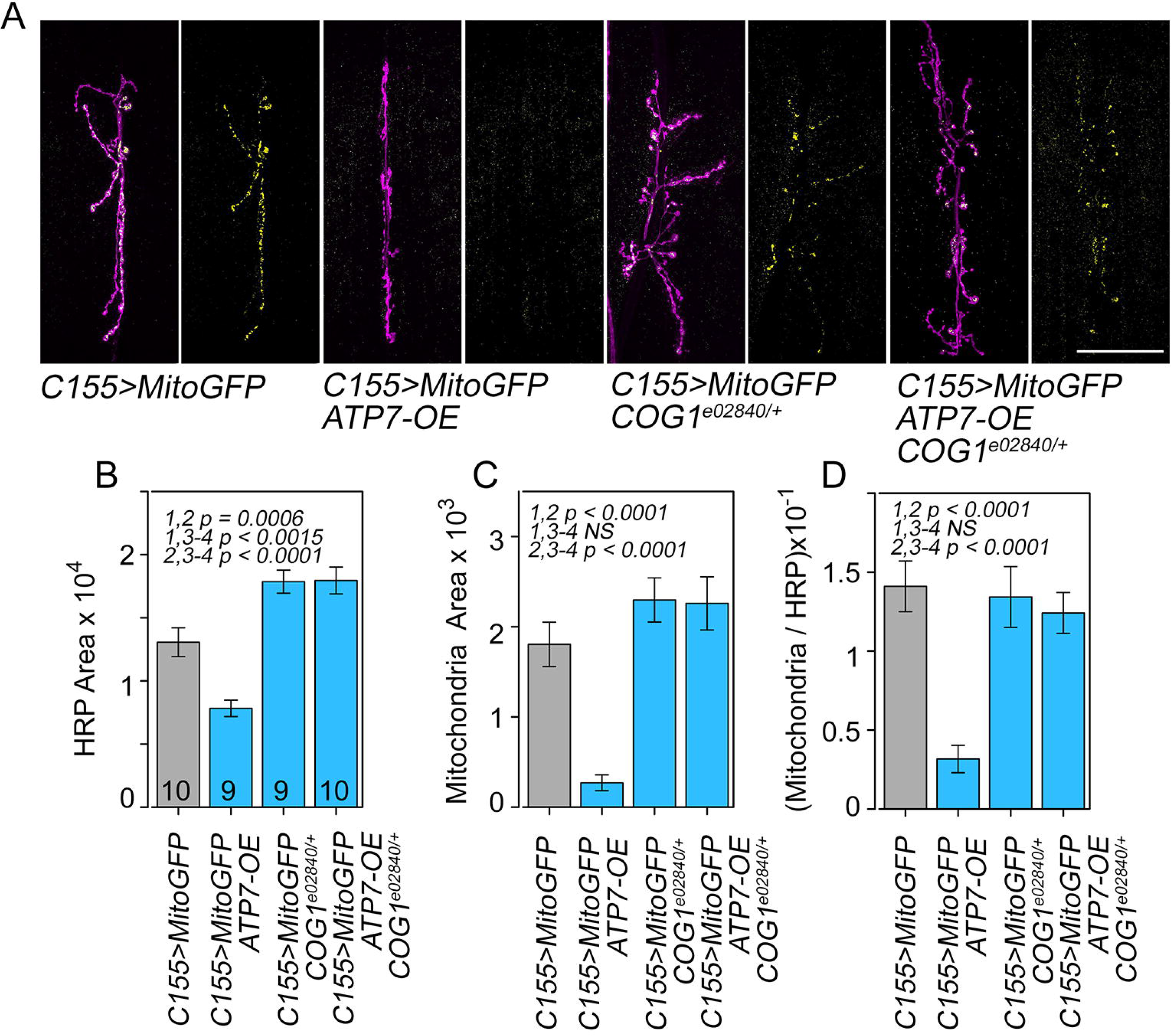
The COG Complex and ATP7 are Required to Maintain Mitochondrial Content in Glutamatergic Synapses. A and B) Muscle 6-7 neuromuscular junctions from third-instar larvae from wild type animals and animals expressing ATP7 transgene, carrying the *COG1*^*e02840/+*^ allele or the combination were crossed to the mitochondrial UAS-mitochondrial-GFP reporter. Reporter genes were expressed with the neuronal C155-GAL4 driver. Dissected larvae were stained with anti-HRP antibodies and anti-GFP Bar 50 μm. B-D) Quantification of synaptic branches and mitochondria content expressed as HRP or GFP area and their ratios. Average ± SEM. n of animals per genotype are presented by the number at the base each column. One Way ANOVA followed by two-tailed Fisher’s Least Significant Difference Comparison. See Extended Data Fig. 6-1

Our results indicate that a single copy loss of the COG1 gene is not sufficient to alter mitochondrial content and distribution in neurons (Fig. 6A-D) even when the COG1 haploinsufficiency increased synapse branching (*COG1*^*e02840/+*^, Fig. 6A and B). In contrast, ATP7 overexpression in neurons resulted in a nearly complete depletion of mitochondria from synapses (Fig. 6A and C-D) with a concomitant increase in mitochondrial signal at ventral nerve cords, a site where motor neuron cell bodies reside (Fig. 6-1). This pronounced mitochondrial synapse depletion phenotype was rescued in animals where the UAS-ATP7 transgene is expressed in the *COG1*^*e02840/+*^ background (Fig. 6A and C-D). The *COG1*^*e02840/+*^-dependent rescue of the ATP7 overexpression mitochondrial phenotype also resulted in a normalization of the mitochondrial signal in the ventral nerve cord (Fig. 6-1).

In parallel, we conducted analyses in *Drosophila* third-instar larvae C-IV da neurons by overexpressing ATP7A, the plasma membrane marker CD4-tdTomato, and MitoGFP in a cell-autonomous manner using the 477- and ppk-GAL4 drivers (Fig. 7). ATP7 overexpression/copper efflux decreased the mitochondrial content in dendrites (Fig. 7A-C). This phenotype was partially rescued by the *COG1*^*e02840/+*^ allele (Fig. 7A), as indicated by a restoration of the mitochondria numbers only at branch points of the dendritic tree (Fig. 7C). These results demonstrate that ATP7, COG, and their interactions modulate dendrite and presynaptic mitochondrial content in motor and sensory neurons. These ATP7 and COG-dependent synaptic branching and mitochondrial phenotypes predict intricate alterations in synaptic transmission and plasticity in neurons with copper dyshomeostasis.

**Fig. 7.**
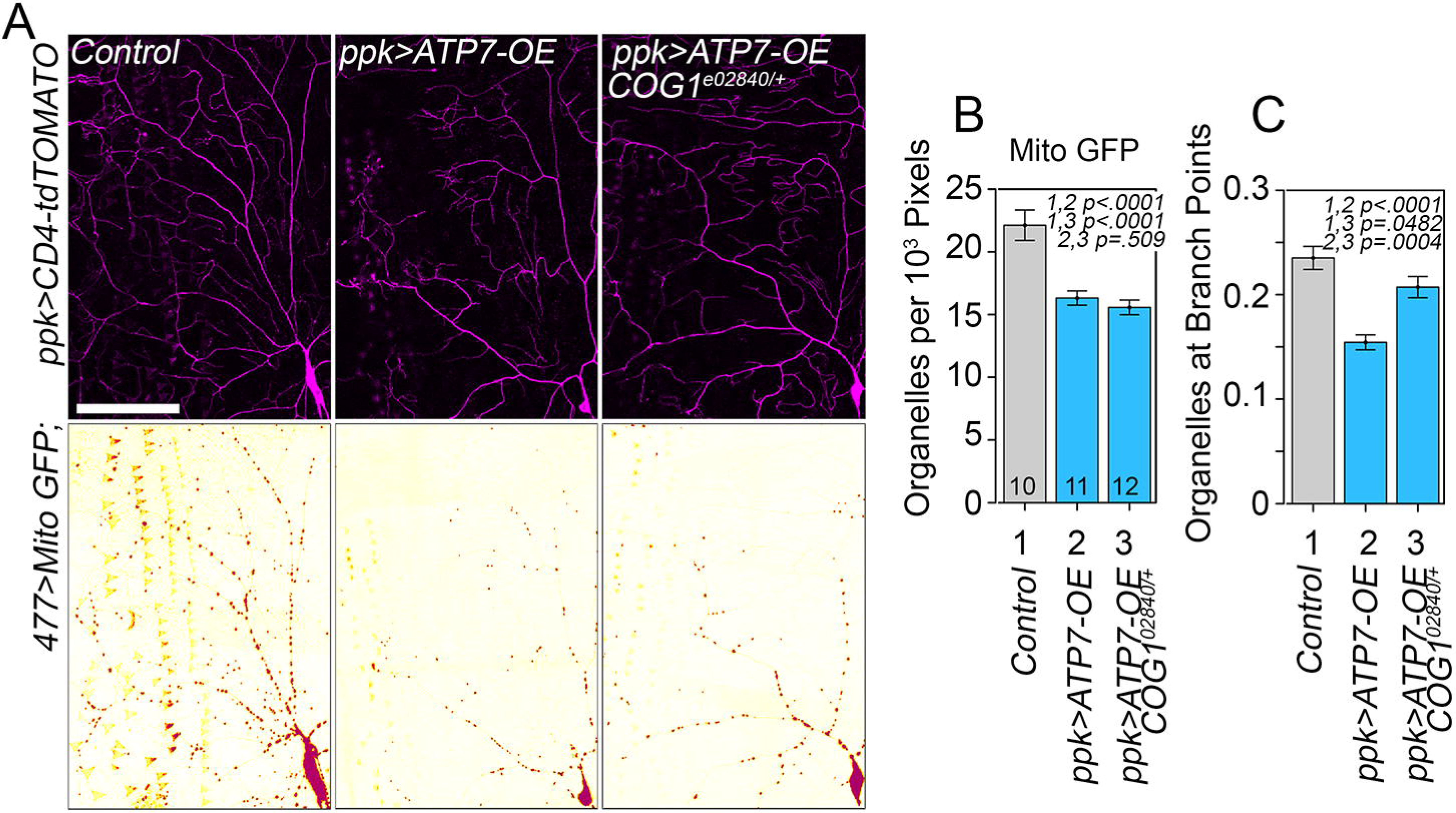
The COG complex and ATP7 are Required to Maintain Mitochondria in Sensory C-IV da Neuron Dendrites. A) Representative live confocal images of C-IV da neurons of the specified genotypes expressing a mitochondria-targeted GFP UAS-transgene and a UAS-plasma membrane marker (CD4-td-Tomato) under the control of the 477-GAL4 driver. B-C show quantification of mitochondria content as organelles per branch length unit or as organelles at branching points. 100 μm. Average ± SEM. n of animals per genotype are presented by the number at the base each column. One Way ANOVA followed by two-tailed Fisher’s Least Significant Difference Comparison.

### The COG complex and ATP7 Modulate Basal Release Probability

We analyzed the effects of single or combined changes in the expression of ATP7 and the COG complex with the *COG1*^*e02840/+*^ allele on neurotransmission (Figs. 8–9). We measured synaptic transmission at the neuromuscular junction of *Drosophila* larvae synapses on muscles 6-7 under two-electrode voltage-clamp. We recorded nerve evoked junctional currents (nEJCs) in the different genotypes at low extracellular Ca^2+^, a condition where basal changes in release probability are easily distinguishable (Acharya et al., 2006; Jorquera et al., 2012; Stevens et al., 2012). Control strains carrying the neuronal GAL4 driver (elav-C155) or the UAS-ATP7 transgene were similar in their nEJCs amplitude (Fig. 8A). However, *COG1*^*e02840/+*^ haploinsufficiency and ATP7 neuronal overexpression/copper efflux displayed increased nEJC amplitudes (Fig. 8A). The nEJCs were larger significantly in COG1 haploinsufficiency and ATP7 neuronal overexpression by 3- and 4-fold, respectively (Fig. 8A-B). This nEJC amplitude phenotype was rescued in *COG1*^*e02840/+*^ animals overexpressing ATP7 in neurons (Fig. 8A-B). The phenotype of large amplitude in nEJCs in *COG1*^*e02840/+*^ animals suggests an increased release probability, as synaptic size and bouton number directly correlate with the amplitude of the overall synaptic responses in healthy synapses (Zhang et al., 2001; Menon et al., 2013). However, in pathological conditions, diverse homeostatic mechanisms of compensation can modify the release probability (Wondolowski and Dickman, 2013; Davis and Muller, 2015; Frank et al., 2020), evident in ATP7A overexpressing neurons, where the amplitude inversely correlates with the synaptic growth phenotype (Fig. 3D, E, and Fig. 8A, B). A synaptic response can be prolonged with increases in release probability, therefore we looked at the nEJCs kinetics. Normalizing average responses during nerve stimulation at low-frequency (Fig. 8C) and estimating a skew values index of kurtosis dividing the charge transferred by their peak amplitude (Fig. 8D), show the slower timing in COG1-haploinsufficiency and ATP7 neuronal overexpression by 2.5ms and 2ms, respectively. This phenotype is rescued partially in animals carrying *COG1*^*e02840/+*^ and expressing the UAS-ATP7 transgene in neurons, indicating that the ATP7 levels modulate amplitude and kinetics of synaptic responses. The neurotransmission phenotype in *COG1*^*e02840/+*^ and neuronal expressed ATP7 animals agree with increased release probability, we further tested this hypothesis measuring facilitation using paired-pulse and tetanic stimulation. To determine the facilitation index in these genotypes, we applied paired-pulse stimulation spaced by 50ms (Fig. 8E-F). Control strains facilitate, *COG1*^*e02840/+*^, and ATP7 neuronal overexpression facilitate similarly (Fig. 8E-F). However, tetanic facilitation after a brief train of stimuli at 20Hz is decreased significantly in animals overexpressing ATP7 in neurons (Fig. 8G-H), indicating a quick saturation in the release probability during nerve activity. These phenotypes were rescued by COG1 haploinsufficiency with the *COG1*^*e02840/+*^ allele (Fig. 8G-H). Altogether, our results demonstrate that ATP7 levels in motor neurons increase the overall initial release probability.

**Fig. 8.**
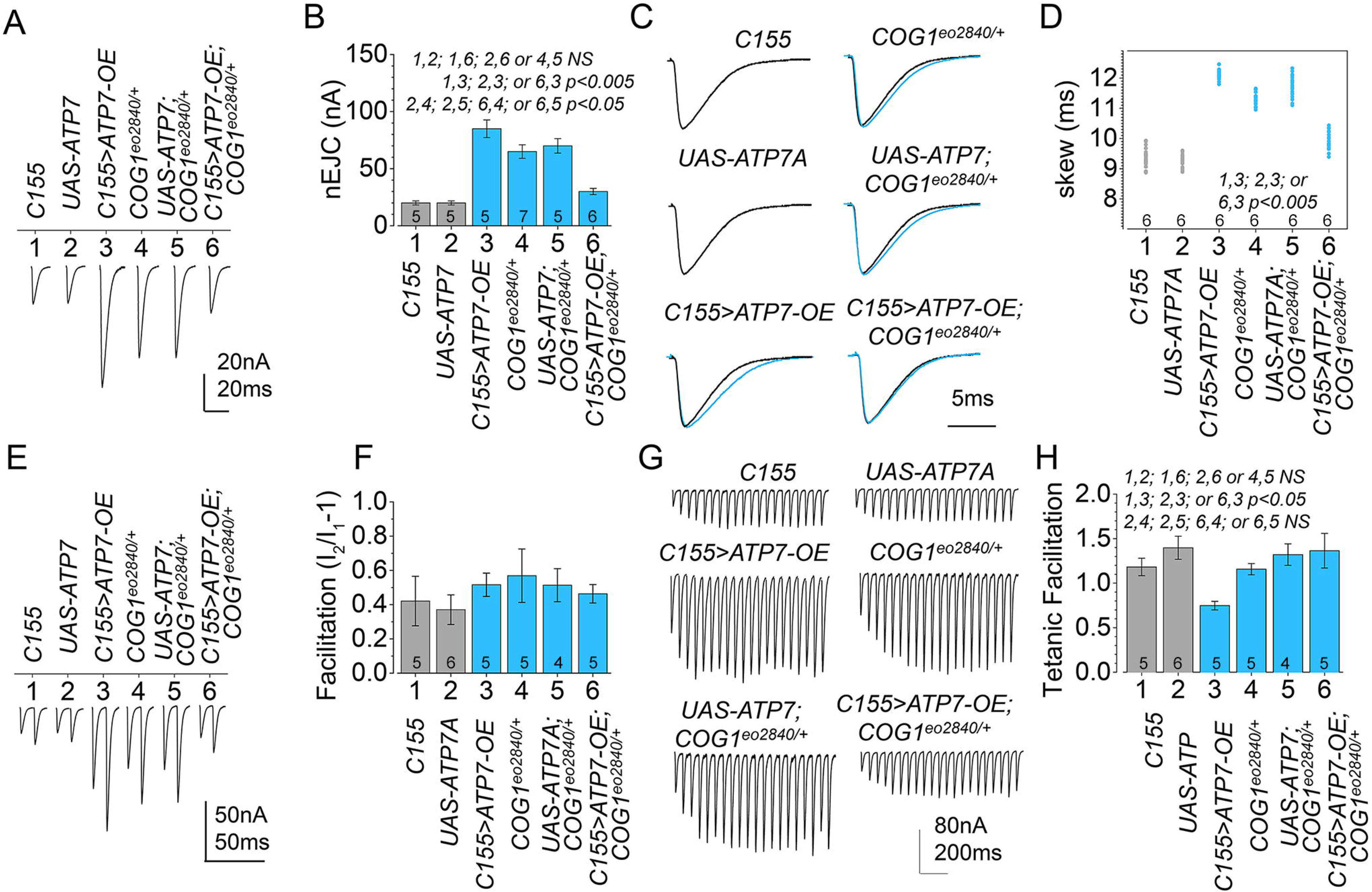
The COG Complex and ATP7 Modifies Nerve-Evoked Synaptic Transmission and Fast Plasticity. A) Representative nEJCs in each genotype at 0.2mM extracellular Ca^2+^ and B) the average amplitude. C) Overlapping of the average normalized nEJCs in each genotype magnified in a new time window and D) the skew factor of kurtosis in each genotype. E) Facilitation in each genotype induced by two pulses spaced at 50ms and F) the average. G) Representative recording of nEJCs during short-term Tetanic Facilitation induced with nerve stimulation at 20Hz, and H) the average tetanic facilitation. n of animals per genotype are presented by the number at the base each column. One Way ANOVA followed by Tukey HSD Post-hoc Test.

**Fig. 9.**
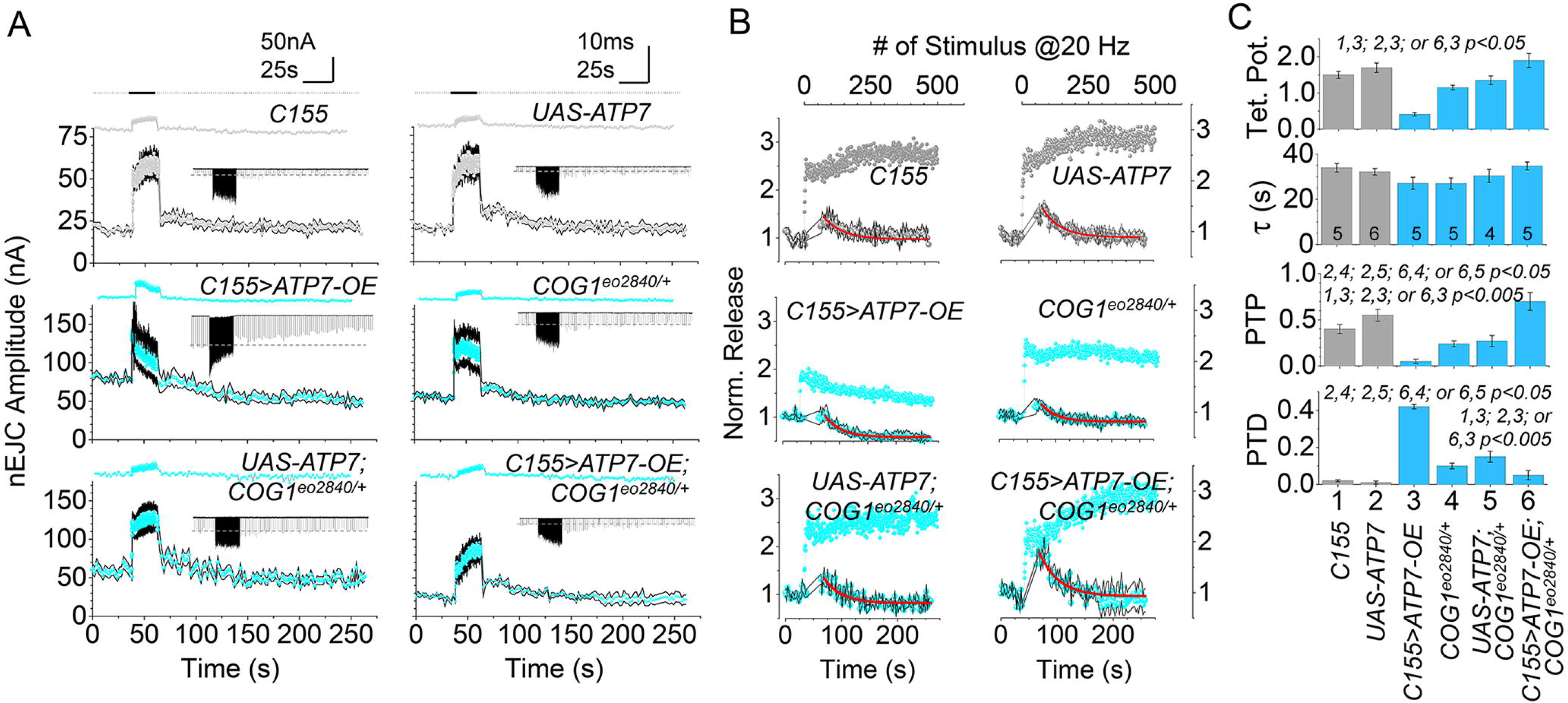
The COG Complex and ATP7 Alter Short-Term Synaptic-Plasticity in Conditioning and Post Conditioning Episodes. A) Average amplitude of nEJCs during the application of a short-term synaptic memory protocol in each genotype, and representative electrophysiological recordings of synaptic transmission (inset). The average instantaneous skewness in each condition is displayed at the top. Test and conditioning stimulation were conducted at 0.5 and 20Hz respectively. B) Average responses during the low-frequency test stimulation (0.5Hz) before and after the tetanic episode normalized by the pre-tetanic value, and the average increase in synaptic efficacy during tetanic activity in each genotype. C) The average estimation for tetanic potentiation, post-tetanic potentiation, and depression, with the time constant of transition. All graphs in c have the same n of animals per genotype. These are presented by the number at the base the tau columns. One Way ANOVA followed by Tukey HSD Post-hoc Test.

### The COG Complex and ATP7 Modify Tetanic and Post-Tetanic Short-Term Synaptic Plasticity

Neuronal transmission is plastic and undergoes short-term changes in synaptic efficacy during high-frequency nerve activation. After the conditioning episode ceases, these changes in synaptic efficacy persist and are progressively restored to their pre-conditioned values over time, a phenomenon known as short-term synaptic memory. Short-term synaptic memory is key for learning and working memory, as determined in *Drosophila* mutants with altered Pavlovian memory retention where neuromuscular synaptic modifications correlate with memory retention phenotypes (Zhong and Wu, 1991). To scrutinize the effects of tetanic and post-tetanic plasticity, we analyzed the nEJCs during a protocol that induces post-tetanic potentiation under low release probability conditions (Fig. 9). In all genotypes, test stimulation for 40 sec at 0.5Hz evoked stable nEJCs (Fig. 9A, B). However, the tetanic episode, induced by 25s of nerve stimulation at 20Hz, dramatically enhance the synaptic transmission decaying in control to the pre-tetanic average values within 50s of test stimulation after the conditioning episode ceased (Fig. 9A, B). However, *COG1*^*e02840/+*^ and neuronal overexpressed ATP7 synapses cannot sustain this increase (Fig. 9A, B). Besides, synaptic transmission decayed after a tetanic episode below the test control in both genotypes, suggesting the presence of a tetanic- and post-tetanic depression. The phenotype observed in synapses overexpressing ATP7 was rescued by the *COG1*^*e02840/+*^ allele (Fig. 9A, B) indicating that the genetic interaction between ATP7 and the COG complex is necessary to sustain short-term plasticity.

To evaluate the presynaptic sources that could modify neurotransmission in these genotypes, we tested the capacity to change their synaptic efficacy by looking in detail at the tetanic and post-tetanic plasticity parameters (Fig. 9C). Normalized tetanic responses indicate that all genotypes display different profiles of enhancement after the initial tetanic facilitation phase (Fig. 9B). Nerve activity induces the characteristic face of augmentation within 10s upon it reaches the maximal potentiation in synaptic enhancement in control strains. Synapses from animals carrying the *COG1*^*e02840/+*^ allele took less time during the augmentation face, also reaching lower potentiation levels (Fig. 9B, C). In turn, ATP7 neuronal overexpression rapidly progressed to depression (Fig. 9A, B, C). This data is consistent with an increase in release probability and abnormal synaptic vesicle availability during periods of high demand. Interestingly, this synaptic behavior resembles the phenotype observed in *Drosophila drp1*, mutants devoid of mitochondria in pre-synapses (Verstreken et al., 2005). However, the synaptic transmission behavior occurring during post-conditioning episodes, such as short-term synaptic memory, is undetermined in this kind of synapses (Verstreken et al., 2005). Pre- and post-tetanic response analysis indicate that control strains induced the characteristic post-tetanic potentiation decaying to pre-tetanic values with a time constant of 45sec (Fig. 9A, B). However, the *COG1*^*e02840/+*^ allele displayed less post-tetanic potentiation which progressed to post tetanic depression (Fig. 9). This post-tetanic potentiation phenotype was even more pronounced in ATP7 overexpressing neurons, which supports the concept of abnormal synaptic vesicle availability during high demand activity. Importantly, these post-tetanic potentiation phenotypes observed in either *COG1*^*e02840/+*^ or ATP7 overexpressing animals were rescued in animals simultaneously carrying the *COG1*^*e02840/+*^ allele and expressing ATP7 in neurons (Fig. 9). Our data demonstrate that genetic interactions between ATP7 and the COG complex, that maintain neuronal copper homeostasis, are necessary for synaptic function, including basal probability of vesicle release, synaptic vesicle availability, and short-term plasticity during conditioning and post-conditioning episodes.

## Discussion

We demonstrate phylogenetically conserved genetic interactions between ATP7 paralogs and a Golgi complex tether, the COG complex. Disruption of these interactions alters presynaptic function, mitochondrial content, and plasticity. *Drosophila* phenotypes caused by changes in the expression of ATP7 are modulated by downregulation of COG complex subunits in diverse neurons. These cell types are affected in Menkes disease by mechanisms yet to be fully explored involving mitochondria (Zlatic et al., 2015; Hartwig et al., 2019; Guthrie et al., 2020). We postulate that genetic and biochemical interactions between ATP7 and COG are necessary to maintain cellular copper homeostasis, which in turn is necessary for mitochondria functional integrity. Our findings support the concept that the Golgi complex and mitochondria are coupled through copper, possibly an elemental ‘second-like messenger’ previously suggested in signaling (Brady et al., 2014; Krishnamoorthy et al., 2016). A Golgi-mitochondria copper dependent communication is also supported by genetic evidence in mammals where mutations in factors required for copper delivery to mitochondria alter the expression of ATP7A and CTR1 (Hlynialuk et al., 2015; Baker et al., 2017).

ATP7A, ATP7B, and CTR1 expression require the COG complex in mammals. The ablation of the COG complex downregulates these copper transporters in mammalian cells in a cell-autonomous manner (Comstra et al., 2017). This observation led us to design a set of genetic tests of copper homeostasis predictions in *Drosophila* tissues. We scrutinized these copper homeostasis predictions using both systemic and cell-autonomous changes in the expression of ATP7, CTR1, and COG complex subunits (Fig. 2). We used ATP7 cell-autonomous overexpression genetic manipulation to induce copper efflux from cells, thus mimicking Menkes-like states. Conversely, ATP7 RNAi was used to increase cellular copper. These are established paradigms to study copper homeostasis in *Drosophila* (Norgate et al., 2006; Burke et al., 2008; Hwang et al., 2014; Mercer et al., 2016). We conclude that ATP7- and CTR1-dependent copper homeostasis is under control of the COG complex in neurons. The following genetic evidence supports this assertion: 1) the rescue of diverse cell-autonomous ATP7 overexpression/copper efflux phenotypes in neuronal tissues by COG gene defects (Figs. 2E, 4A, and 5–9). 2) The COG-dependent rescue of cell-autonomous ATP7 RNAi/copper influx phenotypes in neurons which we infer represents a reduced CTR transport activity at the plasma membrane (Figs. 2G and 4B). 3) The rescue of copper induced mortality phenotype by cell-autonomous downregulation of three COG complex subunits in dopaminergic neurons (Fig. 4B). 4) The rescue of a synaptic haploinsufficiency COG phenotype by systemic overexpression of ATP7 (Fig. 2C). 5) The similar rescue of cell-autonomous ATP7 RNAi/copper influx synaptic phenotypes granted by either neuronal-specific COG subunit or CTR1 downregulation (Fig. 2G). 6) Finally, the rescue of mitochondrial respiration defects by copper ionophores in COG-null human cells (Fig. 5).

While our genetic and biochemical analyses point to a contribution of copper in COG deficient phenotypes in flies and mammalian cells, it is reasonable to expect the contribution of other receptors and membrane transporter to neuronal phenotypes described here. COG genetic defects characteristically impair the activity of glycosyltransferases localized to the Golgi complex (Ungar et al., 2002; Zolov and Lupashin, 2005; Climer et al., 2015; Bailey Blackburn et al., 2016; Blackburn et al., 2018; D’Souza et al., 2019). These enzymes are no longer retained in the Golgi apparatus of COG deficient cells, leading to a global decrease in the glycosylation of proteins traversing the secretory pathway. This defective glycosylation destabilizes membrane proteins residing in the endomembrane system (Ungar et al., 2002; Zolov and Lupashin, 2005; Shestakova et al., 2006; Pokrovskaya et al., 2011; Climer et al., 2015; Bailey Blackburn et al., 2016). These COG-dependent biochemical defects would determine phenotypes in human COG mutations, also known as congenital disorders of glycosylation type II. However, which neuronal proteins undergo defective glycosylation and destabilizations in this disorder is still unclear (Barone et al., 2014; Climer et al., 2015; Climer et al., 2018). We speculate that neurotrophic factors and receptors, synaptic adhesion molecules, calcium and ion channels, and/or synaptic membrane proteins may contribute to the synaptic phenotypes observed in COG deficient *Drosophila* neurons (Kadmon et al., 1990; Han et al., 2004; Kleene and Schachner, 2004; Dong et al., 2008; Kwon and Chapman, 2012; Weiss et al., 2013; Lazniewska and Weiss, 2017).

A prominent phenotype observed in neurons where copper efflux is increased by ATP7 overexpression, and a focus of our studies, is a depletion of mitochondria at neuromuscular synapses and dendrites. This mitochondrial phenotype is concomitant with an accumulation of mitochondria in the ventral nerve cord where motor neuron cell bodies reside. These *Drosophila* mitochondrial phenotypes are reminiscent of mitochondrial defects in Menkes disease (Yoshimura and Kudo, 1983; Onaga et al., 1987; Yamano et al., 1988; Zlatic et al., 2015). We attribute the mitochondrial phenotype in ATP7 overexpressing neurons to defective copper loading in mitochondria, yet this idea needs a direct experimental test as the one performed in a mouse Menkes model (Guthrie et al., 2020). Our observation that COG-null human cells are defective in cellular copper and mitochondrial respiration, the later rescued by copper ionophores, support a model of copper depletion-dependent damage of mitochondria (Fig. 5). Irrespective of whether mitochondria are impaired solely by copper-dependent mechanisms or other additional mechanisms that require copper transporter activity, such as oxidative stress; we postulate that functional mitochondria defects may either prevent their delivery to nerve terminals by impairing mitochondria attachment to kinesin motors, prevent their anchoring to nerve terminals, and/or induce their engulfment by mitophagy, thus trapping them from cell bodies. Copper-dependent damage of mitochondria alters their morphology, uncouples the respiratory chain depolarizing mitochondria, and induces mitophagy (Yoshimura and Kudo, 1983; Yamano and Suzuki, 1985; Onaga et al., 1987; Yamano et al., 1988; Gu et al., 2000; Zischka et al., 2011; Bhattacharjee et al., 2016; Lichtmannegger et al., 2016; Polishchuk et al., 2019). Mitochondrial uncoupling and depolarization change mitochondria kinesin-dependent movement and induce mitophagy in cell bodies (Overly et al., 1996; Miller and Sheetz, 2004; Roberts et al., 2008; Verburg and Hollenbeck, 2008; Wang et al., 2011; Cai et al., 2012; Sheng and Cai, 2012; Ashrafi et al., 2014; Mishra and Chan, 2016; Misgeld and Schwarz, 2017). This evidence suggests that these mitochondrial distribution mechanisms may be engaged in neurons whose copper homeostasis is disrupted by defects in ATP7 and/or the COG complex.

Synapse morphology, mitochondrial content, evoked neurotransmission, and short-term plasticity are modified by changes in the expression of ATP7 and COG complex subunits (Figs. 2–3, 6–9). How these phenotypes are intertwined into cause and effect relationships remains to be explored. We speculate, however, that the absence of mitochondria may account in part for the modified neurotransmission observed in ATP7 overexpression/copper efflux synapses. Mitochondria could contribute to synapses: their calcium buffering capacity (Vos et al., 2010), their ATP synthase activity (Rangaraju et al., 2014), their Krebs cycle metabolites capable of modulating neurotransmission (Ugur et al., 2017), or enzymatic activities necessary for glutamate metabolism (Kovacevic and McGivan, 1983). In addition, mitochondria could offer metabolic flexibility diversifying sources of carbon entering the Krebs cycle for energy production at the synapse (pyruvate, glutamate, or fatty acids) (Olson et al., 2016). As an example, synaptic calcium regulation by mitochondria may explain some of the neurotransmission phenotypes in ATP7 overexpression/copper efflux brains. Short episodes of high-frequency nerve-stimulation induce plasticity in the form of facilitation, augmentation, and potentiation. These are all short-term increases in the efficacy of synaptic transmission, also known as synaptic enhancement (Zucker and Regehr, 2002). Facilitation depends on the residual Ca^2+^ after influx; in turn, augmentation relies on the management of the resting Ca^2+^ during nerve activity, which promotes the exocytosis of the readily releasable pool of vesicles, whereas potentiation depends mostly on the exo/endocytic balance (Zucker et al., 1991; Kamiya and Zucker, 1994; Regehr et al., 1994). Post-tetanic facilitation and augmentation phenomena are related to the decay of residual calcium (Edelman et al., 1987; Zucker et al., 1991; Zucker and Regehr, 2002). However, the resting calcium explains the post tetanic potentiation, indicating that during activity the rising calcium drives phosphorylation-dependent signaling activation, which lasts longer than the calcium transient itself (Macleod et al., 2002; Lnenicka et al., 2006). In that sense, mitochondria could be recruited during the development of post-tetanic potentiation (Tang and Zucker, 1997; Zhong et al., 2001). Thus, mitochondria could maintain or modulate calcium levels at the nerve terminal to sustain diverse forms of synaptic plasticity by buffering the tetanic activity critical for postconditioning episodes like short-term synaptic memory. Our data show that tetanic and post tetanic synaptic enhancements are altered in ATP7 overexpressing/copper efflux synapses, a phenotype that can be reverted by COG complex downregulation. We do not have direct evidence that mitochondria are solely responsible for the synaptic phenotypes in ATP7 overexpression. However, the parallel rescue of both mitochondrial content and neurotransmission phenotypes in ATP7 overexpression synapses after COG complex downregulation support the hypothesis of a mitochondrial-dependent mechanisms account for the neurotransmission phenotypes in these terminals. We propose that copper dyshomeostasis synaptic phenotypes in the fly provide a foundation to understand behavioral and neurological phenotypes characteristic of Menkes and Wilson disease.

## Acknowledgements

This work was supported by grants from the National Institutes of Health 1RF1AG060285 to VF, R01NS108778 to RJ, R15AR070505 to AVM, TelethonTIGEM-CBDM9 to RP, R01NS086082 to DNC, R01GM083144 to VL, and 5K12GM000680-19 to CH. We are indebted to the Faundez lab members for their comments. Stocks obtained from the Bloomington Drosophila Stock Center (NIH P40OD018537) were used in this study.

## Extended Data

**Fig. 4-1.**
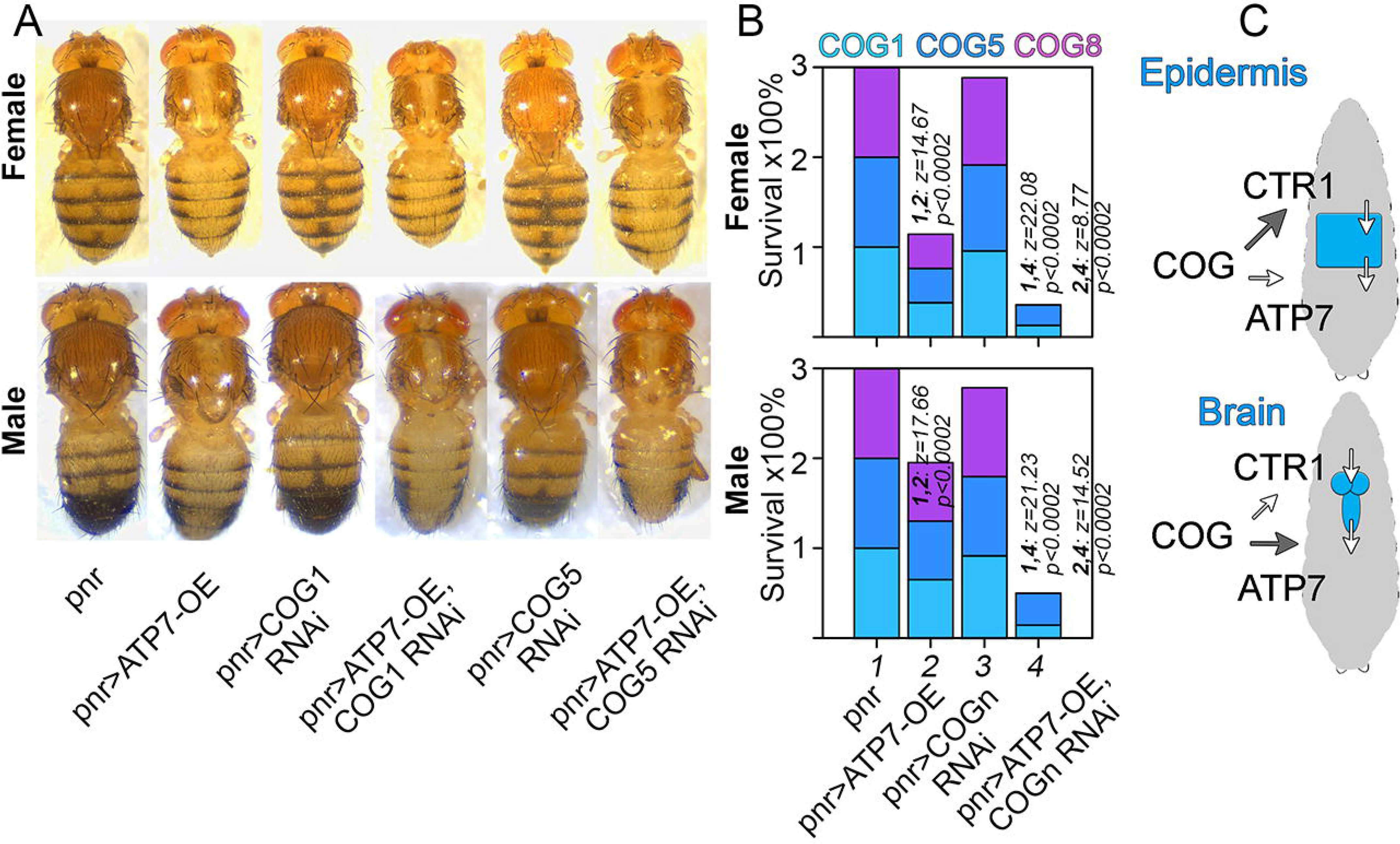
Interactions between ATP7 and the COG Complex are Conserved in *Drosophila* Adult Epidermis. A) Phenotype of transgenic animals expressing UAS-ATP7 or COG subunit RNAi, alone or their combinations, under the control of pnr-GAL4. B) Mortality of adult transgenic animals expressing UAS-ATP7, UAS-COG1, −COG5, or −COG8 RNAi alone or their combinations using the epidermal pnr-GAL4 driver. N=498 total number of animals used in these experiments. p values and z-scores were estimated as two-tailed differences between two proportions. C) Model of the preferential genetic interactions between the COG complex and copper transporters in epidermis or neurons. Arrow size indicates the relative effect of the COG complex in the activity of the copper transporter.

**Fig. 6-1.**
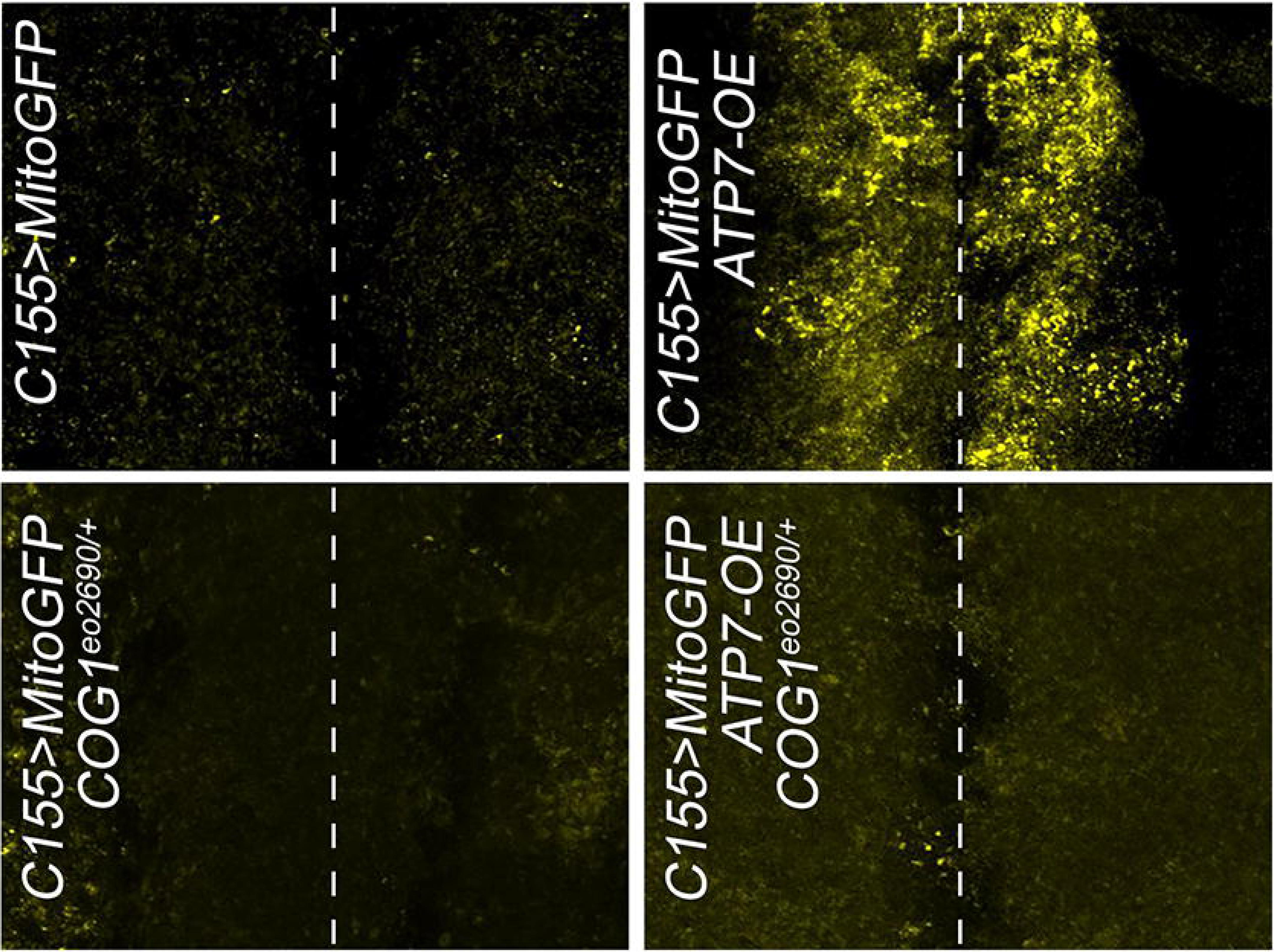
The COG Complex and ATP7 are Required to Maintain Mitochondrial Distribution in Neurons. Third-instar larvae from wild type animals and animals expressing ATP7 transgene, carrying the *COG1*^*e02840/+*^ allele, or their combination were crossed to the mitochondrial UAS-GFP reporter. Transgenes were expressed with the neuronal C155-GAL4 driver. Dissected ventral cords were stained with anti-GFP and imaged by confocal microscopy.

## References

Acharya U, Edwards MB, Jorquera RA, Silva H, Nagashima K, Labarca P, Acharya JK (2006) Drosophila melanogaster Scramblases modulate synaptic transmission. The Journal of cell biology 173:69–82.

Allensworth JL, Evans MK, Bertucci F, Aldrich AJ, Festa RA, Finetti P, Ueno NT, Safi R, McDonnell DP, Thiele DJ, Van Laere S, Devi GR (2015) Disulfiram (DSF) acts as a copper ionophore to induce copper-dependent oxidative stress and mediate anti-tumor efficacy in inflammatory breast cancer. Mol Oncol 9:1155–1168.

Ashrafi G, Schlehe JS, LaVoie MJ, Schwarz TL (2014) Mitophagy of damaged mitochondria occurs locally in distal neuronal axons and requires PINK1 and Parkin. The Journal of cell biology 206:655–670.

Astorga C, Jorquera RA, Ramirez M, Kohler A, Lopez E, Delgado R, Cordova A, Olguin P, Sierralta J (2016) Presynaptic DLG regulates synaptic function through the localization of voltage-activated Ca(2+) Channels. Sci Rep 6:32132.

Bahadorani S, Bahadorani P, Marcon E, Walker DW, Hilliker AJ (2010) A Drosophila model of Menkes disease reveals a role for DmATP7 in copper absorption and neurodevelopment. Disease models & mechanisms 3:84–91.

Bailey Blackburn J, Pokrovskaya I, Fisher P, Ungar D, Lupashin VV (2016) COG Complex Complexities: Detailed Characterization of a Complete Set of HEK293T Cells Lacking Individual COG Subunits. Front Cell Dev Biol 4:23.

Baker ZN, Jett K, Boulet A, Hossain A, Cobine PA, Kim BE, El Zawily AM, Lee L, Tibbits GF, Petris MJ, Leary SC (2017) The mitochondrial metallochaperone SCO1 maintains CTR1 at the plasma membrane to preserve copper homeostasis in the murine heart. Human molecular genetics 26:4617–4628.

Barone R, Fiumara A, Jaeken J (2014) Congenital disorders of glycosylation with emphasis on cerebellar involvement. Semin Neurol 34:357–366.

Berridge MV, Herst PM, Tan AS (2005) Tetrazolium dyes as tools in cell biology: new insights into their cellular reduction. Biotechnol Annu Rev 11:127–152.

Bhattacharjee A, Yang H, Duffy M, Robinson E, Conrad-Antoville A, Lu YW, Capps T, Braiterman L, Wolfgang M, Murphy MP, Yi L, Kaler SG, Lutsenko S, Ralle M (2016) The Activity of Menkes Disease Protein ATP7A Is Essential for Redox Balance in Mitochondria. The Journal of biological chemistry 291:16644–16658.

Binks T, Lye JC, Camakaris J, Burke R (2010) Tissue-specific interplay between copper uptake and efflux in Drosophila. Journal of biological inorganic chemistry: JBIC: a publication of the Society of Biological Inorganic Chemistry 15:621–628.

Blackburn JB, Lupashin VV (2016) Creating Knockouts of Conserved Oligomeric Golgi Complex Subunits Using CRISPR-Mediated Gene Editing Paired with a Selection Strategy Based on Glycosylation Defects Associated with Impaired COG Complex Function. Methods Mol Biol 1496:145–161.

Blackburn JB, Kudlyk T, Pokrovskaya I, Lupashin VV (2018) More than just sugars: Conserved oligomeric Golgi complex deficiency causes glycosylation-independent cellular defects. Traffic 19:463–480.

Brady DC, Crowe MS, Turski ML, Hobbs GA, Yao X, Chaikuad A, Knapp S, Xiao K, Campbell SL, Thiele DJ, Counter CM (2014) Copper is required for oncogenic BRAF signalling and tumorigenesis. Nature 509:492–496.

Burke R, Commons E, Camakaris J (2008) Expression and localisation of the essential copper transporter DmATP7 in Drosophila neuronal and intestinal tissues. Int J Biochem Cell Biol 40:1850–1860.

Cai Q, Zakaria HM, Simone A, Sheng ZH (2012) Spatial parkin translocation and degradation of damaged mitochondria via mitophagy in live cortical neurons. Current biology: CB 22:545–552.

Camakaris J, Danks DM, Ackland L, Cartwright E, Borger P, Cotton RG (1980) Altered copper metabolism in cultured cells from human Menkes’ syndrome and mottled mouse mutants. Biochem Genet 18:117–131.

Clark NL, Alani E, Aquadro CF (2012) Evolutionary rate covariation reveals shared functionality and coexpression of genes. Genome research 22:714–720.

Clark NL, Alani E, Aquadro CF (2013) Evolutionary rate covariation in meiotic proteins results from fluctuating evolutionary pressure in yeasts and mammals. Genetics 193:529–538.

Climer LK, Dobretsov M, Lupashin V (2015) Defects in the COG complex and COG-related trafficking regulators affect neuronal Golgi function. Front Neurosci 9:405.

Climer LK, Hendrix RD, Lupashin VV (2018) Conserved Oligomeric Golgi and Neuronal Vesicular Trafficking. Handb Exp Pharmacol 245:227–247.

Comstra HS, McArthy J, Rudin-Rush S, Hartwig C, Gokhale A, Zlatic SA, Blackburn JB, Werner E, Petris M, D’Souza P, Panuwet P, Barr DB, Lupashin V, Vrailas-Mortimer A, Faundez V (2017) The interactome of the copper transporter ATP7A belongs to a network of neurodevelopmental and neurodegeneration factors. Elife 6.

D’Souza Z, Blackburn JB, Kudlyk T, Pokrovskaya ID, Lupashin VV (2019) Defects in COG-Mediated Golgi Trafficking Alter Endo-Lysosomal System in Human Cells. Front Cell Dev Biol 7:118.

Das R, Bhattacharjee S, Patel AA, Harris JM, Bhattacharya S, Letcher JM, Clark SG, Nanda S, Iyer EPR, Ascoli GA, Cox DN (2017) Dendritic Cytoskeletal Architecture Is Modulated by Combinatorial Transcriptional Regulation in Drosophila melanogaster. Genetics 207:1401–1421.

Davis GW, Muller M (2015) Homeostatic control of presynaptic neurotransmitter release. Annu Rev Physiol 77:251–270.

Divakaruni AS, Paradyse A, Ferrick DA, Murphy AN, Jastroch M (2014) Analysis and interpretation of microplate-based oxygen consumption and pH data. Methods in enzymology 547:309–354.

Dong M, Liu H, Tepp WH, Johnson EA, Janz R, Chapman ER (2008) Glycosylated SV2A and SV2B mediate the entry of botulinum neurotoxin E into neurons. Molecular biology of the cell 19:5226–5237.

Edelman GM, Gall WE, Cowan WM, Neurosciences Institute (New York N.Y.) (1987) Synaptic function. In: The Neurosciences Institute publication series, pp x, 789 p. New York: Wiley.

Findlay GD, Sitnik JL, Wang W, Aquadro CF, Clark NL, Wolfner MF (2014) Evolutionary rate covariation identifies new members of a protein network required for Drosophila melanogaster female post-mating responses. PLoS genetics 10:e1004108.

Foulquier F (2009) COG defects, birth and rise! Biochimica et biophysica acta 1792:896–902.

Frank CA, James TD, Muller M (2020) Homeostatic control of Drosophila neuromuscular junction function. Synapse 74:e22133.

Gokhale A, Vrailas-Mortimer A, Larimore J, Comstra HS, Zlatic SA, Werner E, Manvich DF, Iuvone PM, Weinshenker D, Faundez V (2015a) Neuronal copper homeostasis susceptibility by genetic defects in dysbindin, a schizophrenia susceptibility factor. Human molecular genetics 24:5512–5523.

Gokhale A, Mullin AP, Zlatic S, Easley CA, Merritt ME, Raj N, Larimore J, Gordon DE, Peden AA, Sanyal S, Faundez V (2015b) The N-Ethylmaleimide Sensitive Factor (NSF) and Dysbindin Interact to Modulate Synaptic Plasticity. J Neurosci 35:7643–7653.

Gokhale A, Hartwig C, Freeman AH, Das R, Zlatic SA, Vistein R, Burch A, Carrot G, Lewis AF, Nelms S, Dickman D, Puthenveedu M, Cox DN, Faundez V (2016) The Proteome of BLOC-1 Genetic Defects Identifies the Arp2/3 Actin Polymerization Complex to Function Downstream of the Schizophrenia Susceptibility Factor Dysbindin at the Synapse. J Neurosci 36:12393–12411.

Gu M, Cooper JM, Butler P, Walker AP, Mistry PK, Dooley JS, Schapira AH (2000) Oxidative-phosphorylation defects in liver of patients with Wilson’s disease. Lancet 356:469–474.

Guthrie LM, Soma S, Yuan S, Silva A, Zulkifli M, Snavely TC, Greene HF, Nunez E, Lynch B, De Ville C, Shanbhag V, Lopez FR, Acharya A, Petris MJ, Kim BE, Gohil VM, Sacchettini JC (2020) Elesclomol alleviates Menkes pathology and mortality by escorting Cu to cuproenzymes in mice. Science 368:620–625.

Han W, Rhee JS, Maximov A, Lao Y, Mashimo T, Rosenmund C, Sudhof TC (2004) N-glycosylation is essential for vesicular targeting of synaptotagmin 1. Neuron 41:85–99.

Hartwig C, Zlatic SA, Wallin M, Vrailas-Mortimer A, Fahrni CJ, Faundez V (2019) Trafficking mechanisms of P-type ATPase copper transporters. Current opinion in cell biology 59:24–33.

Hlynialuk CJ, Ling B, Baker ZN, Cobine PA, Yu LD, Boulet A, Wai T, Hossain A, El Zawily AM, McFie PJ, Stone SJ, Diaz F, Moraes CT, Viswanathan D, Petris MJ, Leary SC (2015) The Mitochondrial Metallochaperone SCO1 Is Required to Sustain Expression of the High-Affinity Copper Transporter CTR1 and Preserve Copper Homeostasis. Cell reports 10:933–943.

Horn D, Barrientos A (2008) Mitochondrial copper metabolism and delivery to cytochrome c oxidase. IUBMB Life 60:421–429.

Hwang JE, de Bruyne M, Warr CG, Burke R (2014) Copper overload and deficiency both adversely affect the central nervous system of Drosophila. Metallomics: integrated biometal science 6:2223–2229.

Iyer EP, Iyer SC, Sullivan L, Wang D, Meduri R, Graybeal LL, Cox DN (2013a) Functional genomic analyses of two morphologically distinct classes of Drosophila sensory neurons: post-mitotic roles of transcription factors in dendritic patterning. PloS one 8:e72434.

Iyer SC, Ramachandran Iyer EP, Meduri R, Rubaharan M, Kuntimaddi A, Karamsetty M, Cox DN (2013b) Cut, via CrebA, transcriptionally regulates the COPII secretory pathway to direct dendrite development in Drosophila. Journal of cell science 126:4732–4745.

Jorquera RA, Huntwork-Rodriguez S, Akbergenova Y, Cho RW, Littleton JT (2012) Complexin controls spontaneous and evoked neurotransmitter release by regulating the timing and properties of synaptotagmin activity. J Neurosci 32:18234–18245.

Kadmon G, Kowitz A, Altevogt P, Schachner M (1990) Functional cooperation between the neural adhesion molecules L1 and N-CAM is carbohydrate dependent. The Journal of cell biology 110:209–218.

Kaler SG (2011) ATP7A-related copper transport diseases-emerging concepts and future trends. Nature reviews Neurology 7:15–29.

Kamiya H, Zucker RS (1994) Residual Ca2+ and short-term synaptic plasticity. Nature 371:603–606.

Kleene R, Schachner M (2004) Glycans and neural cell interactions. Nat Rev Neurosci 5:195–208.

Kovacevic Z, McGivan JD (1983) Mitochondrial metabolism of glutamine and glutamate and its physiological significance. Physiological reviews 63:547–605.

Kranz C, Ng BG, Sun L, Sharma V, Eklund EA, Miura Y, Ungar D, Lupashin V, Winkel RD, Cipollo JF, Costello CE, Loh E, Hong W, Freeze HH (2007) COG8 deficiency causes new congenital disorder of glycosylation type IIh. Human molecular genetics 16:731–741.

Krishnamoorthy L, Cotruvo JA, Jr., Chan J, Kaluarachchi H, Muchenditsi A, Pendyala VS, Jia S, Aron AT, Ackerman CM, Wal MN, Guan T, Smaga LP, Farhi SL, New EJ, Lutsenko S, Chang CJ (2016) Copper regulates cyclic-AMP-dependent lipolysis. Nat Chem Biol 12:586–592.

Kuo YM, Zhou B, Cosco D, Gitschier J (2001) The copper transporter CTR1 provides an essential function in mammalian embryonic development. Proceedings of the National Academy of Sciences of the United States of America 98:6836–6841.

Kwon SE, Chapman ER (2012) Glycosylation is dispensable for sorting of synaptotagmin 1 but is critical for targeting of SV2 and synaptophysin to recycling synaptic vesicles. The Journal of biological chemistry 287:35658–35668.

Lazniewska J, Weiss N (2017) Glycosylation of voltage-gated calcium channels in health and disease. Biochim Biophys Acta Biomembr 1859:662–668.

Lee CE, Singleton KS, Wallin M, Faundez V (2020) Rare Genetic Diseases: Nature’s Experiments on Human Development. iScience 23:101123.

Lichtmannegger J et al. (2016) Methanobactin reverses acute liver failure in a rat model of Wilson disease. J Clin Invest 126:2721–2735.

Lnenicka GA, Grizzaffi J, Lee B, Rumpal N (2006) Ca2+ dynamics along identified synaptic terminals in Drosophila larvae. J Neurosci 26:12283–12293.

Lutsenko S, Barnes NL, Bartee MY, Dmitriev OY (2007) Function and regulation of human copper-transporting ATPases. Physiological reviews 87:1011–1046.

Macleod GT, Hegstrom-Wojtowicz M, Charlton MP, Atwood HL (2002) Fast calcium signals in Drosophila motor neuron terminals. J Neurophysiol 88:2659–2663.

McAllum EJ, Hare DJ, Volitakis I, McLean CA, Bush AI, Finkelstein DI, Roberts BR (2020) Regional iron distribution and soluble ferroprotein profiles in the healthy human brain. Progress in neurobiology 186:101744.

Menkes JH (1988) Kinky hair disease: twenty five years later. Brain & development 10:77–79.

Menkes JH (1999) Menkes disease and Wilson disease: two sides of the same copper coin. Part I: Menkes disease. European journal of paediatric neurology: EJPN: official journal of the European Paediatric Neurology Society 3:147–158.

Menon KP, Carrillo RA, Zinn K (2013) Development and plasticity of the Drosophila larval neuromuscular junction. Wiley Interdiscip Rev Dev Biol 2:647–670.

Mercer SW, La Fontaine S, Warr CG, Burke R (2016) Reduced glutathione biosynthesis in Drosophila melanogaster causes neuronal defects linked to copper deficiency. Journal of neurochemistry 137:360–370.

Miller KE, Sheetz MP (2004) Axonal mitochondrial transport and potential are correlated. Journal of cell science 117:2791–2804.

Misgeld T, Schwarz TL (2017) Mitostasis in Neurons: Maintaining Mitochondria in an Extended Cellular Architecture. Neuron 96:651–666.

Mishra P, Chan DC (2016) Metabolic regulation of mitochondrial dynamics. The Journal of cell biology 212:379–387.

Morgan MT, Bourassa D, Harankhedkar S, McCallum AM, Zlatic SA, Calvo JS, Meloni G, Faundez V, Fahrni CJ (2019) Ratiometric two-photon microscopy reveals attomolar copper buffering in normal and Menkes mutant cells. Proceedings of the National Academy of Sciences of the United States of America 116:12167–12172.

Mullin AP, Sadanandappa MK, Ma W, Dickman DK, VijayRaghavan K, Ramaswami M, Sanyal S, Faundez V (2015) Gene Dosage in the Dysbindin Schizophrenia Susceptibility Network Differentially Affect Synaptic Function and Plasticity. J Neurosci 35:325–338.

Norgate M, Lee E, Southon A, Farlow A, Batterham P, Camakaris J, Burke R (2006) Essential roles in development and pigmentation for the Drosophila copper transporter DmATP7. Molecular biology of the cell 17:475–484.

Olson KA, Schell JC, Rutter J (2016) Pyruvate and Metabolic Flexibility: Illuminating a Path Toward Selective Cancer Therapies. Trends Biochem Sci 41:219–230.

Onaga A, Kawasaki H, Yamano T, Shimada M, Nishimura M (1987) Light and electron microscopic study on cerebellar cortex of macular mutant mouse as a model of Menkes kinky hair disease. Brain & development 9:265–269.

Overly CC, Rieff HI, Hollenbeck PJ (1996) Organelle motility and metabolism in axons vs dendrites of cultured hippocampal neurons. Journal of cell science 109 (Pt 5):971–980.

Petris MJ, Mercer JF, Culvenor JG, Lockhart P, Gleeson PA, Camakaris J (1996) Ligand-regulated transport of the Menkes copper P-type ATPase efflux pump from the Golgi apparatus to the plasma membrane: a novel mechanism of regulated trafficking. The EMBO journal 15:6084–6095.

Pokrovskaya ID, Willett R, Smith RD, Morelle W, Kudlyk T, Lupashin VV (2011) Conserved oligomeric Golgi complex specifically regulates the maintenance of Golgi glycosylation machinery. Glycobiology 21:1554–1569.

Polishchuk EV et al. (2019) Activation of autophagy, observed in liver tissues from patients with Wilson disease and from ATP7B-deficient animals, protects hepatocytes from copper-induced apoptosis. Gastroenterology in press.

Polishchuk R, Lutsenko S (2013) Golgi in copper homeostasis: a view from the membrane trafficking field. Histochemistry and cell biology 140:285–295.

Rai D, Dey S, Ray K (2018) A method for estimating relative changes in the synaptic density in Drosophila central nervous system. BMC Neurosci 19:30.

Rangaraju V, Calloway N, Ryan TA (2014) Activity-driven local ATP synthesis is required for synaptic function. Cell 156:825–835.

Regehr WG, Delaney KR, Tank DW (1994) The role of presynaptic calcium in short-term enhancement at the hippocampal mossy fiber synapse. J Neurosci 14:523–537.

Roberts EA, Robinson BH, Yang S (2008) Mitochondrial structure and function in the untreated Jackson toxic milk (tx-j) mouse, a model for Wilson disease. Mol Genet Metab 93:54–65.

Rodriguez A, Ehlenberger DB, Dickstein DL, Hof PR, Wearne SL (2008) Automated three-dimensional detection and shape classification of dendritic spines from fluorescence microscopy images. PloS one 3:e1997.

Ruepp A, Waegele B, Lechner M, Brauner B, Dunger-Kaltenbach I, Fobo G, Frishman G, Montrone C, Mewes HW (2010) CORUM: the comprehensive resource of mammalian protein complexes––2009. Nucleic acids research 38:D497–501.

Sanders SJ et al. (2019) A framework for the investigation of rare genetic disorders in neuropsychiatry. Nat Med 25:1477–1487.

Schindelin J, Arganda-Carreras I, Frise E, Kaynig V, Longair M, Pietzsch T, Preibisch S, Rueden C, Saalfeld S, Schmid B, Tinevez JY, White DJ, Hartenstein V, Eliceiri K, Tomancak P, Cardona A (2012) Fiji: an open-source platform for biological-image analysis. Nat Methods 9:676–682.

Sheng ZH, Cai Q (2012) Mitochondrial transport in neurons: impact on synaptic homeostasis and neurodegeneration. Nat Rev Neurosci 13:77–93.

Shestakova A, Zolov S, Lupashin V (2006) COG complex-mediated recycling of Golgi glycosyltransferases is essential for normal protein glycosylation. Traffic 7:191–204.

Singhania A, Grueber WB (2014) Development of the embryonic and larval peripheral nervous system of Drosophila. Wiley Interdiscip Rev Dev Biol 3:193–210.

Stevens RJ, Akbergenova Y, Jorquera RA, Littleton JT (2012) Abnormal synaptic vesicle biogenesis in Drosophila synaptogyrin mutants. J Neurosci 32:18054–18067, 18067a.

Tang Y, Zucker RS (1997) Mitochondrial involvement in post-tetanic potentiation of synaptic transmission. Neuron 18:483–491.

Ugur B, Bao H, Stawarski M, Duraine LR, Zuo Z, Lin YQ, Neely GG, Macleod GT, Chapman ER, Bellen HJ (2017) The Krebs Cycle Enzyme Isocitrate Dehydrogenase 3A Couples Mitochondrial Metabolism to Synaptic Transmission. Cell reports 21:3794–3806.

Ungar D, Oka T, Brittle EE, Vasile E, Lupashin VV, Chatterton JE, Heuser JE, Krieger M, Waters MG (2002) Characterization of a mammalian Golgi-localized protein complex, COG, that is required for normal Golgi morphology and function. The Journal of cell biology 157:405–415.

Verburg J, Hollenbeck PJ (2008) Mitochondrial membrane potential in axons increases with local nerve growth factor or semaphorin signaling. J Neurosci 28:8306–8315.

Verstreken P, Ly CV, Venken KJ, Koh TW, Zhou Y, Bellen HJ (2005) Synaptic mitochondria are critical for mobilization of reserve pool vesicles at Drosophila neuromuscular junctions. Neuron 47:365–378.

Vos M, Lauwers E, Verstreken P (2010) Synaptic mitochondria in synaptic transmission and organization of vesicle pools in health and disease. Front Synaptic Neurosci 2:139.

Wang X, Winter D, Ashrafi G, Schlehe J, Wong YL, Selkoe D, Rice S, Steen J, LaVoie MJ, Schwarz TL (2011) PINK1 and Parkin target Miro for phosphorylation and degradation to arrest mitochondrial motility. Cell 147:893–906.

Weiss N, Black SA, Bladen C, Chen L, Zamponi GW (2013) Surface expression and function of Cav3.2 T-type calcium channels are controlled by asparagine-linked glycosylation. Pflugers Arch 465:1159–1170.

Willett R, Blackburn JB, Climer L, Pokrovskaya I, Kudlyk T, Wang W, Lupashin V (2016) COG lobe B sub-complex engages v-SNARE GS15 and functions via regulated interaction with lobe A sub-complex. Sci Rep 6:29139.

Wondolowski J, Dickman D (2013) Emerging links between homeostatic synaptic plasticity and neurological disease. Front Cell Neurosci 7:223.

Wu X, Steet RA, Bohorov O, Bakker J, Newell J, Krieger M, Spaapen L, Kornfeld S, Freeze HH (2004) Mutation of the COG complex subunit gene COG7 causes a lethal congenital disorder. Nat Med 10:518–523.

Xiao T, Ackerman CM, Carroll EC, Jia S, Hoagland A, Chan J, Thai B, Liu CS, Isacoff EY, Chang CJ (2018) Copper regulates rest-activity cycles through the locus coeruleus-norepinephrine system. Nat Chem Biol 14:655–663.

Yamano T, Suzuki K (1985) Abnormalities of Purkinje cell arborization in brindled mouse cerebellum. A Golgi study. Journal of neuropathology and experimental neurology 44:85–96.

Yamano T, Shimada M, Onaga A, Kawasaki H, Iwane S, Ono K, Nishimura M (1988) Electron microscopic study on brain of macular mutant mouse after copper therapy. Acta neuropathologica 76:574–580.

Yang CH, Rumpf S, Xiang Y, Gordon MD, Song W, Jan LY, Jan YN (2009) Control of the postmating behavioral switch in Drosophila females by internal sensory neurons. Neuron 61:519–526.

Yoshimura N, Kudo H (1983) Mitochondrial abnormalities in Menkes’ kinky hair disease (MKHD). Electron-microscopic study of the brain from an autopsy case. Acta neuropathologica 59:295–303.

Zhang YQ, Bailey AM, Matthies HJ, Renden RB, Smith MA, Speese SD, Rubin GM, Broadie K (2001) Drosophila fragile X-related gene regulates the MAP1B homolog Futsch to control synaptic structure and function. Cell 107:591–603.

Zhong N, Beaumont V, Zucker RS (2001) Roles for mitochondrial and reverse mode Na+/Ca2+ exchange and the plasmalemma Ca2+ ATPase in post-tetanic potentiation at crayfish neuromuscular junctions. J Neurosci 21:9598–9607.

Zhong Y, Wu CF (1991) Altered synaptic plasticity in Drosophila memory mutants with a defective cyclic AMP cascade. Science 251:198–201.

Zischka H, Lichtmannegger J, Schmitt S, Jagemann N, Schulz S, Wartini D, Jennen L, Rust C, Larochette N, Galluzzi L, Chajes V, Bandow N, Gilles VS, DiSpirito AA, Esposito I, Goettlicher M, Summer KH, Kroemer G (2011) Liver mitochondrial membrane crosslinking and destruction in a rat model of Wilson disease. J Clin Invest 121:1508–1518.

Zlatic S, Comstra HS, Gokhale A, Petris MJ, Faundez V (2015) Molecular basis of neurodegeneration and neurodevelopmental defects in Menkes disease. Neurobiology of disease 81:154–161.

Zlatic SA, Vrailas-Mortimer A, Gokhale A, Carey LJ, Scott E, Burch R, McCall MM, Rudin-Rush S, Davis JB, Hartwig C, Werner E, Li L, Petris M, Faundez V (2018) Rare Disease Mechanisms Identified by Genealogical Proteomics of Copper Homeostasis Mutant Pedigrees. Cell Syst 6:368–380 e366.

Zolov SN, Lupashin VV (2005) Cog3p depletion blocks vesicle-mediated Golgi retrograde trafficking in HeLa cells. The Journal of cell biology 168:747–759.

Zucker RS, Regehr WG (2002) Short-term synaptic plasticity. Annu Rev Physiol 64:355–405.

Zucker RS, Delaney KR, Mulkey R, Tank DW (1991) Presynaptic calcium in transmitter release and posttetanic potentiation. Annals of the New York Academy of Sciences 635:191–207.

